# Synthetic RNA-seq cohorts for data sharing: a discovery-aware benchmark at transcriptome scale

**DOI:** 10.64898/2026.05.22.726357

**Authors:** Aditya Nanda, Somdutta Saha

## Abstract

**Background:** Sharing patient-level gene expression data is essential for translational discovery but carries documented re-identification risks. Bulk RNA-seq count matrices can retain genotypic signals and paired clinical metadata compounds this through quasi-identifier matching. Synthetic RNA-seq cohorts offer a complementary path for privacy-preserving data sharing, but the field lacks a multi-axis benchmark that probes biological fidelity and empirical privacy risk at transcriptome scale. Here we present a multi-axis benchmark framework that reflects how transcriptomic cohorts are used in translational practice.

**Methods:** We benchmarked three generative models across four cohorts drawn from datasets spanning oncology (TCGA-LUAD), sepsis (GSE184900), and pediatric IBD (RISK/GSE57945): `dbTwin` (a non-deep-learning, target-conditioned method that operates natively at RNA-seq scale), `class-MVN` (a low-rank target-conditioned multivariate Gaussian model), and `PCA-CTGAN` (a tabular GAN trained in PCA-compressed space). Synthetic cohorts were generated from training folds of a five-fold stratified design. We evaluated DE genes recovery, log_2_FC and significance (padj) concordance, held-out AUC (TSTR) and SHAP concordance and distance-based memorization risk.

**Results:** class-MVN recovered **64.8**% and **43.1**% of real DE genes in the two binary cohorts with high fold-change correlation but lower significance concordance (*r* = **0.24**–**0.68**) and inflated DE gene counts. `dbTwin` recovered **78.7**% and **91.8**% of real DE genes in the same cohorts, with high fold-change correlation and stronger significance concordance (*r* ≥ **0.88**). Both methods matched held-out real AUC under TSTR, but SHAP agreement differed substantially: `dbTwin` preserved feature attribution patterns across cohorts (SHAP top-50 genes *r* = **0.84**–**0.99** across two binary and two multiclass cohorts), whereas `class-MVN` showed moderate performance for majority classes but degraded in multiclass and imbalanced settings (SHAP *r* = **0.31**–**0.79**). `PCA-CTGAN` performed poorly across most DE and ML metrics. Distance-toclosest-record analysis did not indicate memorization by any of the models.

**Conclusions:** We introduced a multi-axis, transcriptome-scale, discovery-aware benchmark to validate synthetic RNA-seq cohorts for translational workflows and evaluated three generative models across four real-world cohorts. These results support the use of synthetic RNA-seq cohorts for exploratory analysis and method development, while emphasizing the need for careful validation before use in higher-stakes applications. All benchmark code and data are available at https://github.com/Nanda-Aditya/rna-syn-bench.

## Introduction

The Transcriptome occupies a singular position in biology: it functions as the proximate bridge anchoring genotype and natural selection processes on one end and phenotype on the other [1–3]. RNA sequencing has become the standard instrument for transcriptome-wide measurement in disease and health [4], while RNA-seq data has accumulated at unprecedented scale [5, 6].

Although RNA-seq cohorts—gene expression counts paired with clinical metadata—are covered by the Common Rule, patient-level access remains difficult: Data Use Agreements, IRB review, and related governance requirements impose barriers, especially in rare-disease and cross-institution settings [7, 8]. These barriers are compounded by mounting evidence that RNA-seq data is reidentifiable: prior work has mapped risk across the processing stack from BAM/VCF byproducts to allele-specific signals [9–12], and we focus here on bulk counts with matched clinical metadata— the most common format in end-user workflows.

Schadt et al. showed that an eQTL-based Bayesian model can infer individual SNP genotypes directly from expression counts [13]; Harmanci and Gerstein further demonstrated that adding demographic attributes (sex and ethnicity) meaningfully increased the fraction of vulnerable individuals beyond eQTL matching alone [14], a directly relevant finding given that translational RNA-seq cohorts almost always carry matched clinical metadata with such quasi-identifiers. Though eQTL-based prediction was once thought limited by population heterogeneity [15], recent attacks have closed that gap: Walker et al. achieved near-perfect linking-attack accuracy on both single-cell and pseudobulked count matrices [16], and Sadhuka et al. achieved perfect re-identification of all 138 GTEx held-out individuals using counts from just four chromosomes and 135 of 138 using a single chromosome [17].

Predicted genotypes (SNP vectors) are sufficient for re-identification via consumer genealogy or genomic databases: Erlich et al. showed that a database covering just 2% of a target population provides a third-cousin-or-closer match for nearly any member [18]. Edge and Coop cleverly used ∼900 synthetic genomes to GEDmatch and used it as an oracle to recover allele data across 82% of the genome for European-ancestry individuals [19], and Ney et al. reconstructed 92% of a target’s SNP profile with 98% accuracy by exploiting GEDmatch’s visual chromosome-comparison tool [20]. Crucially, phased haplotypes are not required: tools for unphased genotype data reliably detect distant kinship from raw genotype calls alone [21, 22].

Taken together, this literature establishes not merely that re-identification is possible, but that the attack surface is expanding faster and the attack chain from expression to identity is increasingly complete: expression counts leak genotypes, genotypes leak identity, and the reference databases needed to complete the chain are growing. Current policy treats the distinction between raw reads and processed counts as a meaningful privacy boundary when the evidence shows it is not [23]; a scoping review of 42 original studies confirmed that attack feasibility continues to grow as public and breached genetic databases expand [24]. Some authors have characterized the genomic re-identification risks as theoretical rather than current, advocating for governance-based controls over data restriction: Byrd et al. assumed high computational barriers and privileged data access [7], while an ELSI program at the GetPreCiSe Center concluded re-identification risk is “often overstated” [25]. Both assessments have since been substantially weakened by the discriminative and linking attacks described above [16, 17]. History is instructive here. Homer et al. ca. 2008 showed that aggregate GWAS summary statistics, once considered safe, were sufficient for individual membership inference [26], prompting NIH to restrict their open release. Processed RNA-seq counts occupy an analogous position today and the policy environment is shifting accordingly: in December 2025, NIH issued a Request for Information (NOT-OD-26-023) proposing that transcriptomic data (including counts) be designated controlled-access data [27].

Taken together, these developments establish that access controls around transcriptomic data are tightening and that the research community will increasingly require layered technical and policy mechanisms to enable legitimate data sharing as the regulatory landscape evolves. Beyond public repositories, such constraints also impose friction on bilateral collaborations, where Data Use Agreements and IRB review can delay data exchange by months even when both parties have legitimate research intent [7, 8]. These developments highlight the need for complementary approaches to privacy-preserving data sharing that can operate alongside, rather than in place of, access-control frameworks. Against this backdrop, synthetic data has emerged as a candidate solution to the problem of privacy-preserving genomic data sharing and reuse [8, 23]. Synthetic data, broadly defined, is artificially generated records that retain aggregate analytical properties of a real dataset while preserving privacy at the level of individual samples or records. It offers a complementary path: one that does not require the infrastructure commitments of federated learning approaches [28, 29] and produces portable datasets suitable for exploratory analysis and hypothesis generation prior to controlled-access validation on real data.

Although synthetic data itself has a storied history in physics, biology, and engineering [30–34], its broader adoption in healthcare as a privacy-preserving mechanism for mixed-type tabular datasets was catalysed by the development of deep generative models — generative adversarial networks (GANs) and variational autoencoders (VAEs) [35] and diffusion models [36]. A parallel line of work, initiated by synthpop [37], replaces deep-learning architectures with tree-based sequential synthesis (CART, LightGBM, random forest), building records column-by-column [38, 39]; methods in both families have been coupled with formal differential privacy guarantees [40, 41]. Both traditions have been extensively applied to structured healthcare records, with multiple systematic reviews surveying this landscape [42–44].

Clinico-transcriptomic data poses challenges qualitatively distinct from structured EHR or clinical records. Expression matrices are ultra-high-dimensional (≈20,000–50,000 genes per sample) with complex covariance structure from co-expression networks and pathway organisation [5, 45], and clinical annotations must remain jointly coherent with the expression profile, *i.e.*, a synthetic EGFR-mutant sample must recapitulate the characteristic transcriptomic signature or it has no analytical value. Standard tabular generative models neither scale to this feature space nor enforce cross-modal consistency. A benchmark of 11 generative models on RNA-seq cohorts found that most capture basic distributional fidelity but fail on translationally meaningful metrics [46].

In this work, we make two contributions. First, we present a discovery-aware, multi-axis benchmark whose evaluation axes reflect how transcriptomic cohorts are actually used in translational practice, spanning differential expression for hit prioritization, SHAP-based feature attribution for biomarker discovery, held-out ML utility, and empirical privacy risk. Second, we introduce and benchmark dbTwin, a non-deep-learning, target-conditioned generator that operates natively at transcriptome scale, alongside two baseline models spanning statistical (class-MVN) and deeplearning (PCA-CTGAN) approaches. We evaluate all three generators on four clinico-transcriptomic cohorts from three public datasets spanning oncology (TCGA-LUAD), sepsis (GSE184900), and pediatric IBD (RISK/GSE57945). All benchmark and evaluation code is released to enable reproducible validation of any synthetic cohort.

## Results

### Overall benchmarking paradigm

Our benchmark tested synthetic generators across four clinico-transcriptomic cohorts spanning 3 datasets in oncology, pediatric IBD and Sepsis as tabulated in Table 1. We used two binary classification tasks to assess whether synthetic cohorts preserve case–control and mutation-specific expression signals: `LUAD-EGFR` (EGFR mutation status in TCGA lung adenocarcinoma [48, 51]) and `Sepsis` (bloodstream infection case–control from GSE184900 [49]). Separately, two multiclass tasks test preservation of finer-grained biological structure: predicting tumor stage (`Stage I, Stage II` and `Advanced`), using the same TCGA-LUAD cohort and predicting disease classification (`Crohn’s disease / ulcerative colitis / non-IBD` controls from the RISK pediatric inception cohort, GSE57945 [50]). Together, these pairs span distinct tissues (lung tumor, whole blood, ileal biopsy), sample sizes (n ≈ 150 – 500), and classification granularities, providing a diverse testbed for assessing the utility of synthetic cohorts at transcriptome scale.

**Table 1.** Summary of dataset-clinical target pairs used in the benchmark. Each column defines a unique evaluation unit comprising a clinico-transcriptomic cohort paired with a specific prediction target. Short labels are used throughout the text and figures.

|  | LUAD-EGFR | Sepsis | LUAD-Stage | RISK-IBD |
| --- | --- | --- | --- | --- |
| **Source** | TCGA-LUAD [47, 48] | GSE184900[49] | TCGA-LUAD | GSE57945 (RISK) [50] |
| **Tissue** | Lung tumor | Whole blood | Lung tumor | Ileal biopsy |
| **Target** | EGFR mutation-wild-type | Blood Sepsis-control | Tumor stage (I, II and Adv.) | IBD subtype (CD / UC / Control) |
| **Classes** | **2** | **2** | **3** | **3** |
| **N<sub>samples</sub>** | **510** | **152** | **510** | **322** |
| **N<sub>genes</sub>** | **19730** | **27101** | **19730** | **43795** |
| **Task type** | Binary | Binary | Multiclass | Multiclass |

We first established a unified evaluation framework as shown in Fig. 1. Panel A illustrates the data partitioning strategy: each of the four clinico-transcriptomic datasets was split via five-fold stratified sampling into a training set and a held-out test set. The three generators were each fit exclusively on training data and produced synthetic cohorts matching the real data in schema, sample size, and class distribution - yielding one synthetic replicate per fold and five replicates per generator.

**Figure 1.**
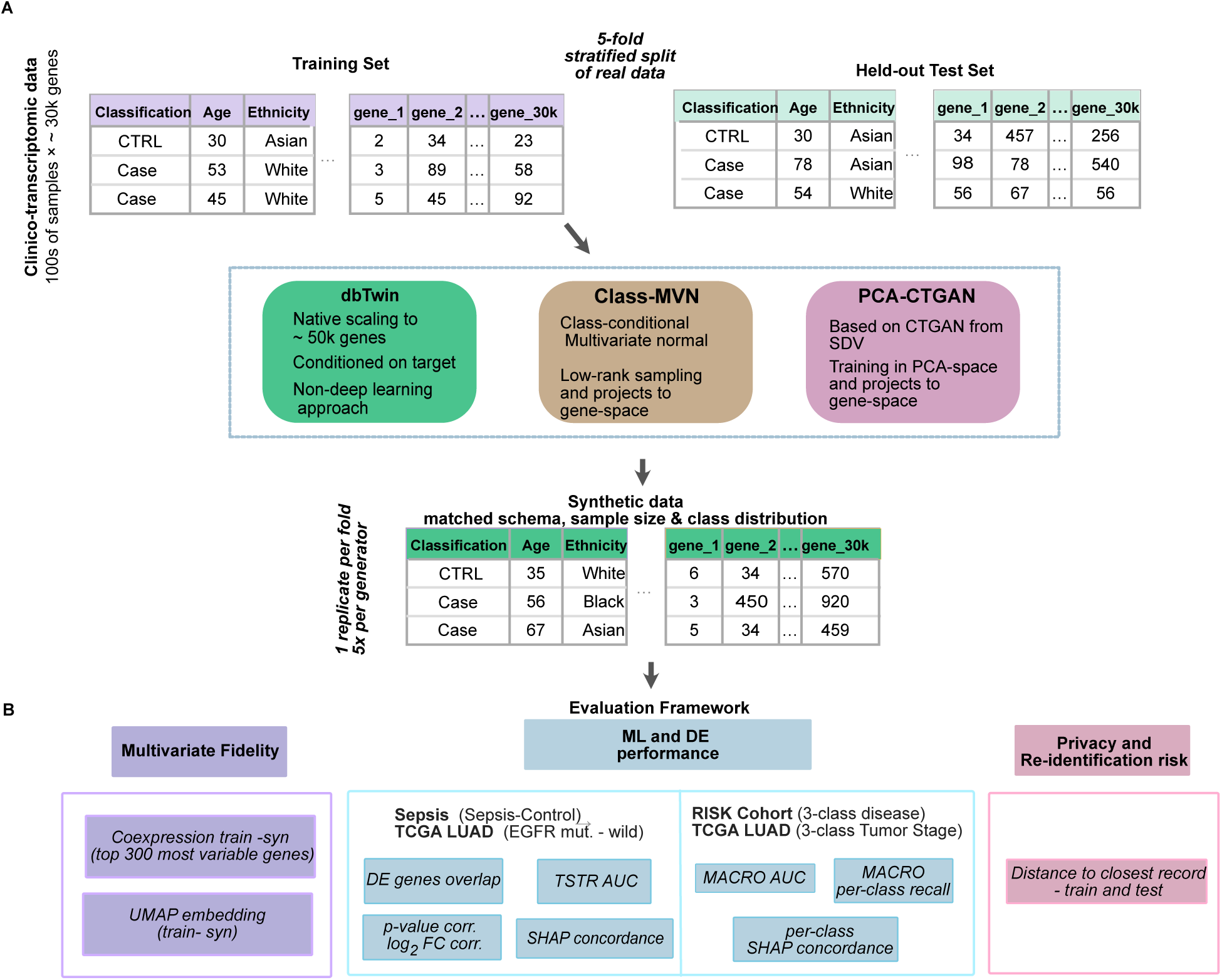
Overview of the synthetic data generation and evaluation framework. (A) Clinico-transcriptomic data 100s of subjects × 30 k genes) are partitioned via five-fold stratified splitting into a *training set* used to fit generative models and a *held-out test set*. Three generators are compared: `dbTwin` (a non-deep-learning, target-conditioned model), `class-MVN` (a class-conditional multivariate normal baseline), and `PCA-CTGAN` (a CTGAN variant operating in PCA space). Each generator produces synthetic cohorts that match the real data in schema, sample size, and class distribution (one replicate per fold, five replicates per generator). **(B)** The evaluation framework spans three axes. *Multivariate fidelity*: co-expression preservation and UMAP embedding overlap, *ML and DE performance*: for binary tasks (`Sepsis` blood case– control; `LUAD-EGFR` mutation), volcano plots (log_2_FC and padj), DE gene overlap, TSTR AUC, and SHAP concordance at gene-level are reported; for multiclass tasks (`RISK-IBD` and `LUAD-Stage`), macro AUC, macro per-class recall, and per-class, gene-level SHAP concordance are reported. *Privacy and re-identification risk*: distance to closest record relative to both train and test sets.

We tested 3 different generators in detail:

- dbTwin: a proprietary non-deep-learning, target-conditioned method natively scaling to 50 000 genes. `dbTwin` extends efficient linear algebra-based algorithms developed for synthetic data generation from large-scale brain activity recordings - functional MRI, Electrocorticography, Calcium imaging [33, 34] - and has been optimized for RNA-seq cohorts.
- class-MVN: a scalable low-rank extension of `class-MVN` (multi-variate normal) proposed in [46].
- PCA-CTGAN: an extension of CTGAN [35] operating in PCA-compressed space.

Panel B details the three evaluation axes as follows:

- Multivariate fidelity assessed using co-expression similarity and UMAP manifold alignment (see Fig. 2).
- Biological utility assessed by using Differential Expression and Machine Learning analyses detailed in Figures 3 – 7.
- Privacy/re-identification risk assessed using standard Distance to closest record between synthetic and real expression values (Fig. 9).

**Figure 2.**
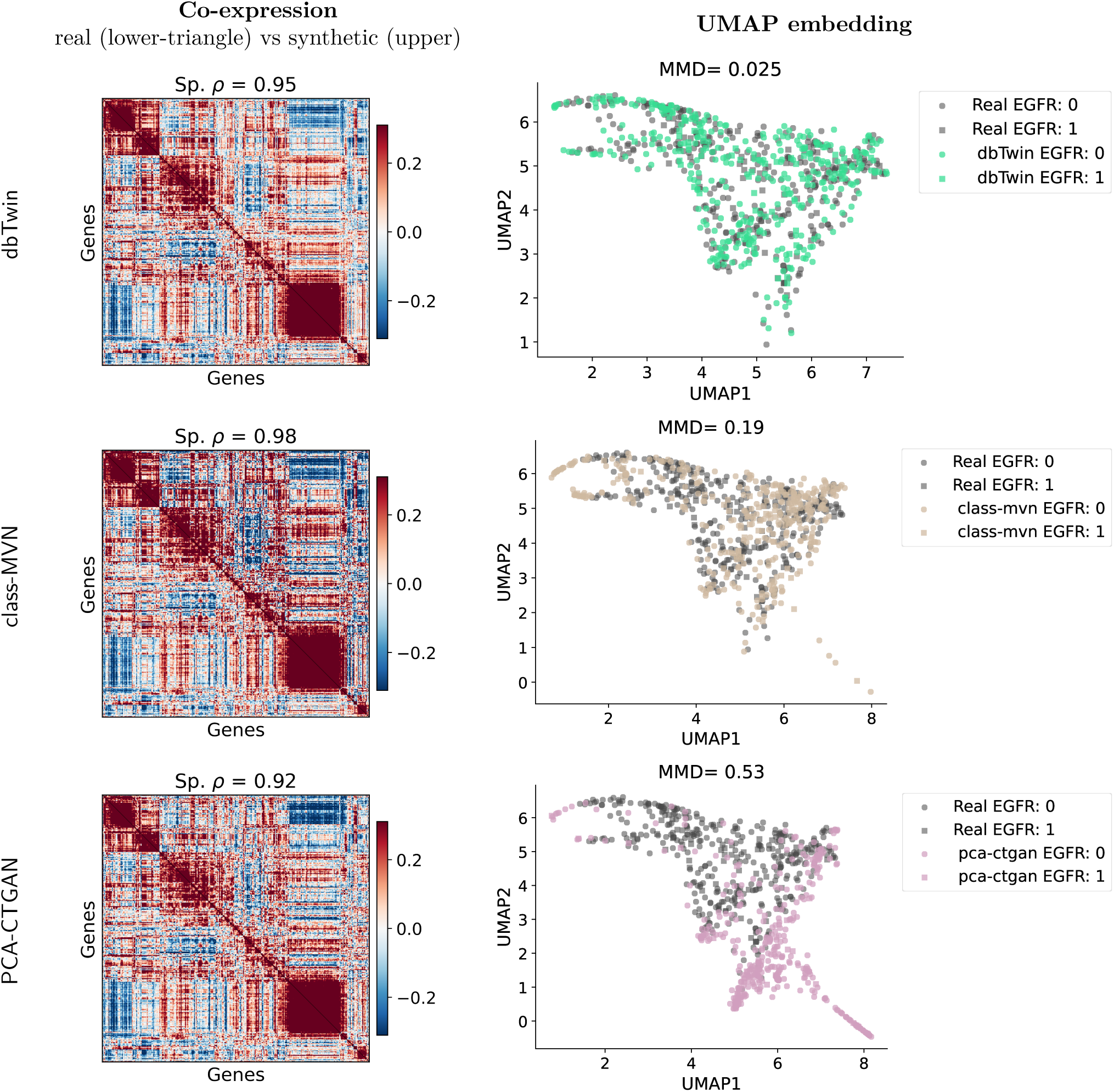
Multivariate fidelity: co-expression and UMAP embedding. Results shown for `LUAD-EGFR` dataset. Please see SI for the remaining 3 cohorts. **Left column:** Gene co-expression correlation matrices comparing real data (lower triangle) versus synthetic data (upper triangle) for the top 300 most variable genes. Hierarchical clustering leaf order is computed from the real data and applied to each synthetic cohort. **Right column:** UMAP embeddings (components 1 vs 2) showing training data (grey points) overlaid with synthetic data (colored points) for each generator.

**Figure 3.**
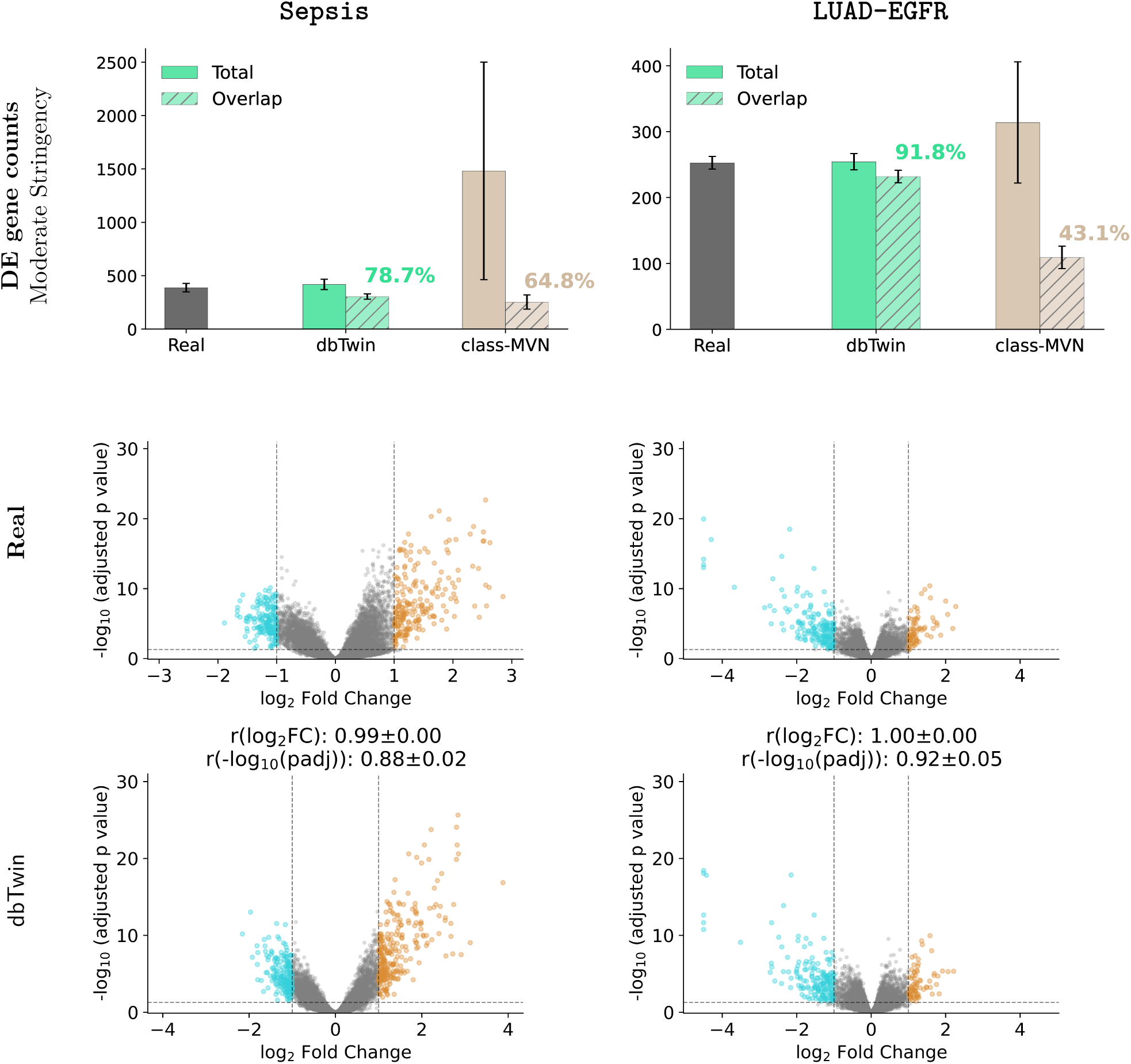
Differential expression fidelity on binary classification tasks (Part 1). Results are shown for the **Sepsis** (blood case–control; left column) and **`LUAD-EGFR`** (right column) datasets. **Row 1:** Number of DE genes detected in the real versus synthetic data and their overlap (moderate stringency: *p*_adj_ *<* 0.05, |log_2_FC| ≥ 1, baseMean ≥ 10; error-bars denote mean ± s.d. over 5 folds). **Rows 2–3:** Volcano plots showing DE significance (− log_10_ adjusted *p*-value) versus effect size (log_2_ fold-change) for real data and `dbTwin` synthetic data. Significant DE genes are highlighted. `PCA-CTGAN` (not-shown) underperformed with less than 10% real DE genes recovery. See Fig. 4 for volcano plots for models: `class-MVN` and `PCA-CTGAN`.

**Figure 4.**
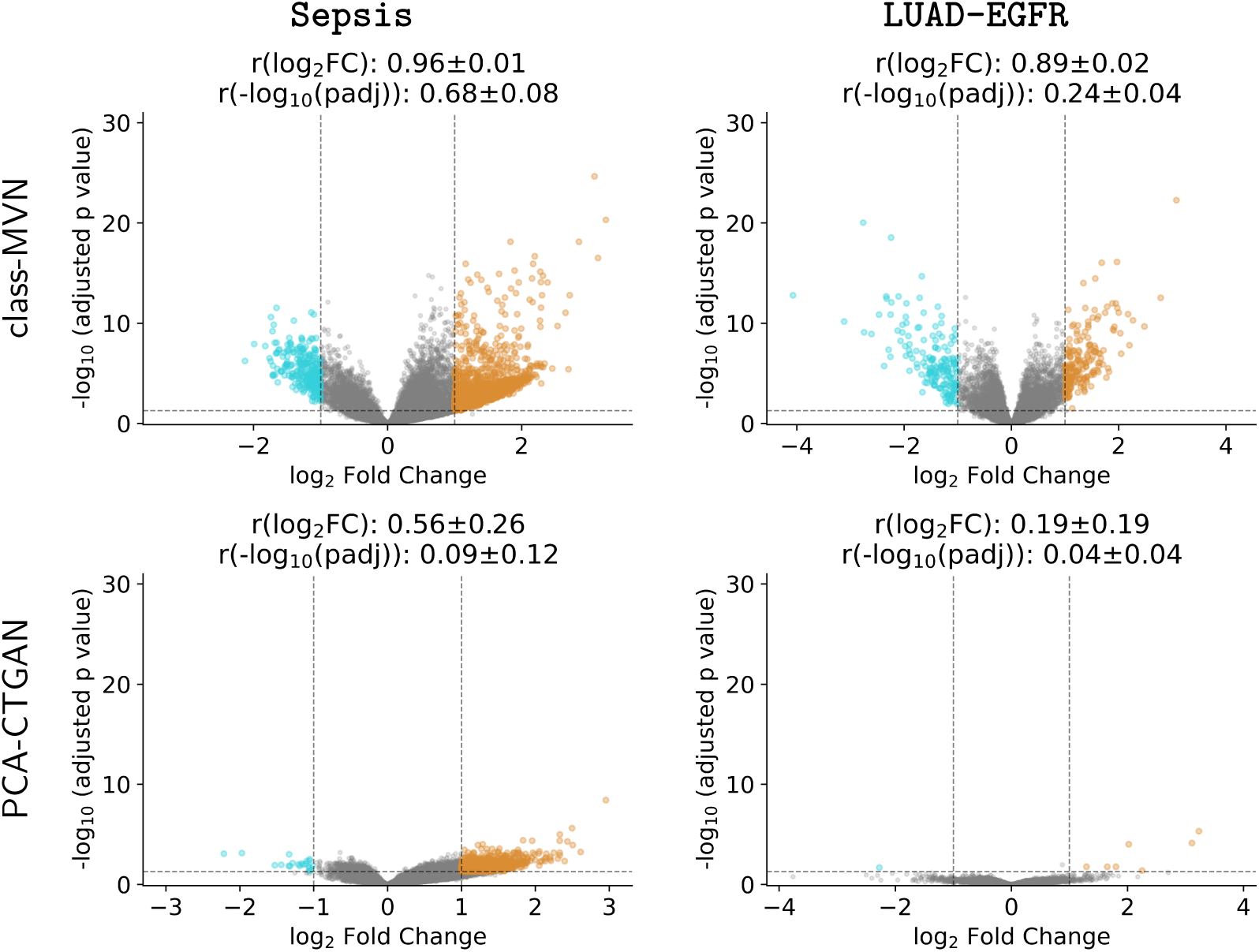
Differential expression fidelity on binary classification tasks (Part 2). Results are shown for the **`Sepsis`** (blood case–control; left column) and **`LUAD-EGFR`** (right column) datasets. Volcano plots showing DE significance (− log_10_ adjusted *p*-value) versus effect size (log_2_ fold-change) for synthetic data from `class-MVN` and `PCA-CTGAN` generators. Significant DE genes (*p*_adj_ *<* 0.05, |log_2_FC| ≥ 1, *baseMean* ≥ 10) are highlighted. See Fig. 3 for DE overlap, real data, and `dbTwin` results.

**Figure 5.**
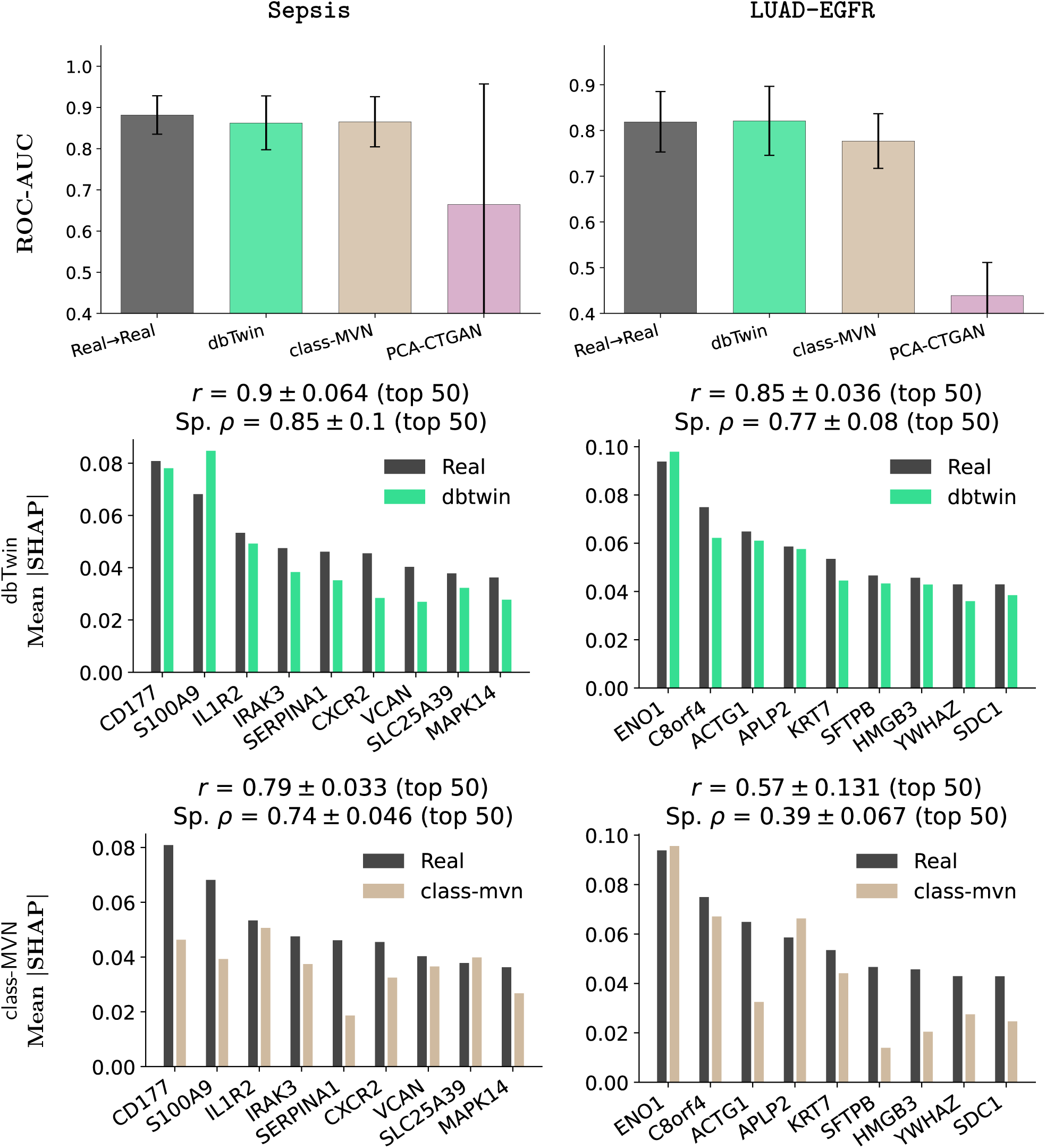
Machine learning utility on binary classification tasks. `Sepsis` (blood case–control; left column) and **`LUAD-EGFR`** (right column) datasets. **Top row:** Held-out test-set AUC for models trained on real data (TRTR, dark bars) versus synthetic data (TSTR, lighter bars). Each bar shows mean ± s.d. over five replicates. A smaller TRTR–TSTR gap indicates higher analytical utility of the synthetic cohort. **Rows 2–3: SHAP gene-feature concordance**: Mean absolute gene-level SHAP weights from classifiers trained on real training (black) versus synthetic data (colored), both evaluated on the held-out real test set. Annotations indicate Pearson correlation (*r*) for the top 50 genes and Spearman rank correlation (Sp. *ρ*) for the top 50 genes (mean ± SD) over 5 folds. Only median-auc-fold is shown for both models: `dbTwin`(left) and `class-MVN`(right). Complete results for all folds are shown in SI.

**Figure 6.**
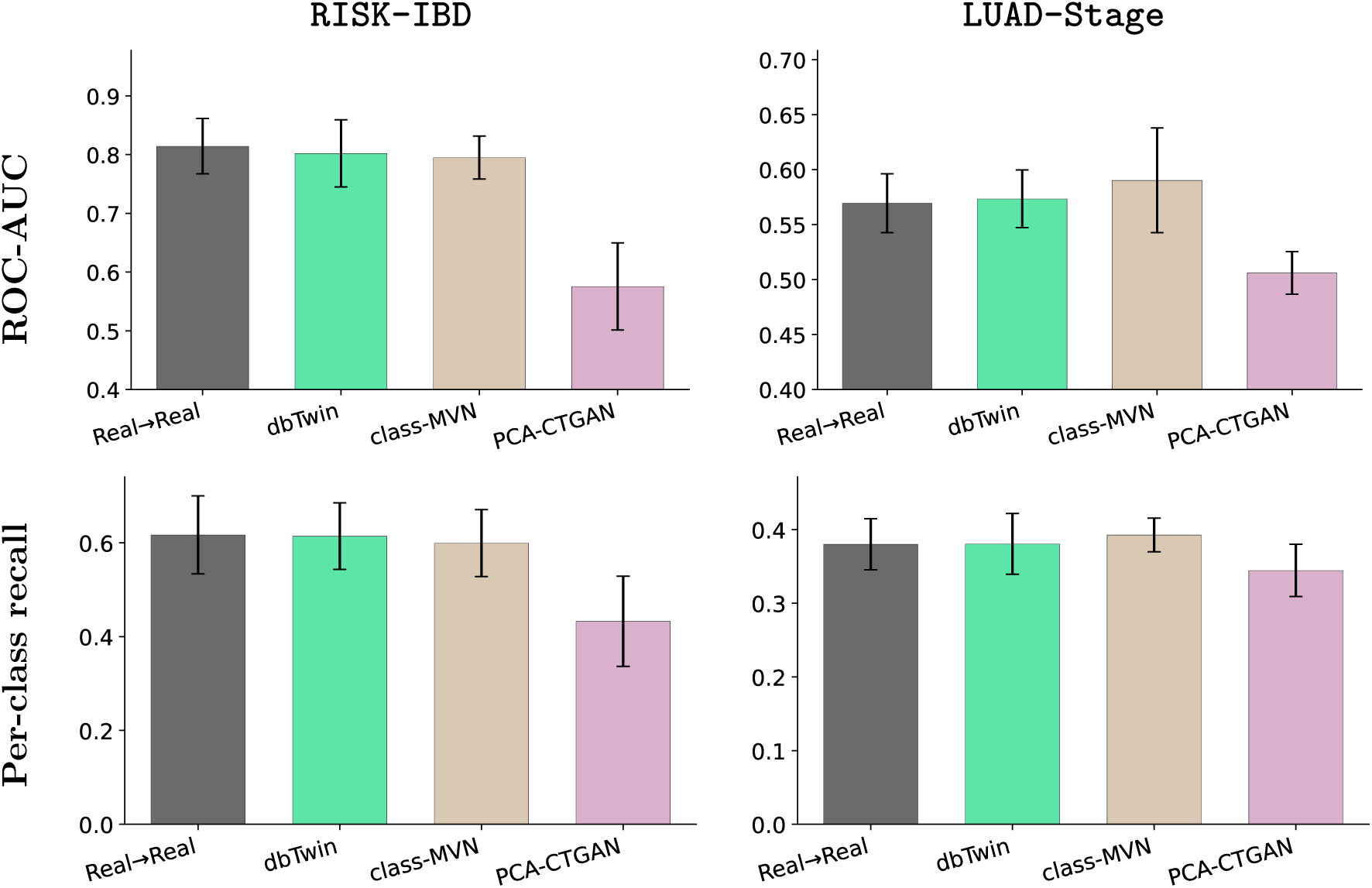
Multiclass ML on RISK-IBD and LUAD-Stage: **Top row:** Held-out test-set macro-averaged AUC for models trained on real data (TRTR, dark bars) versus synthetic data (TSTR, lighter bars). **Bottom row:** Macro-averaged per-class recall (PCR) for models trained on real data (TRTR, dark bars) versus synthetic data (TSTR, lighter bars) showing balanced accuracy across classes.

**Figure 7.**
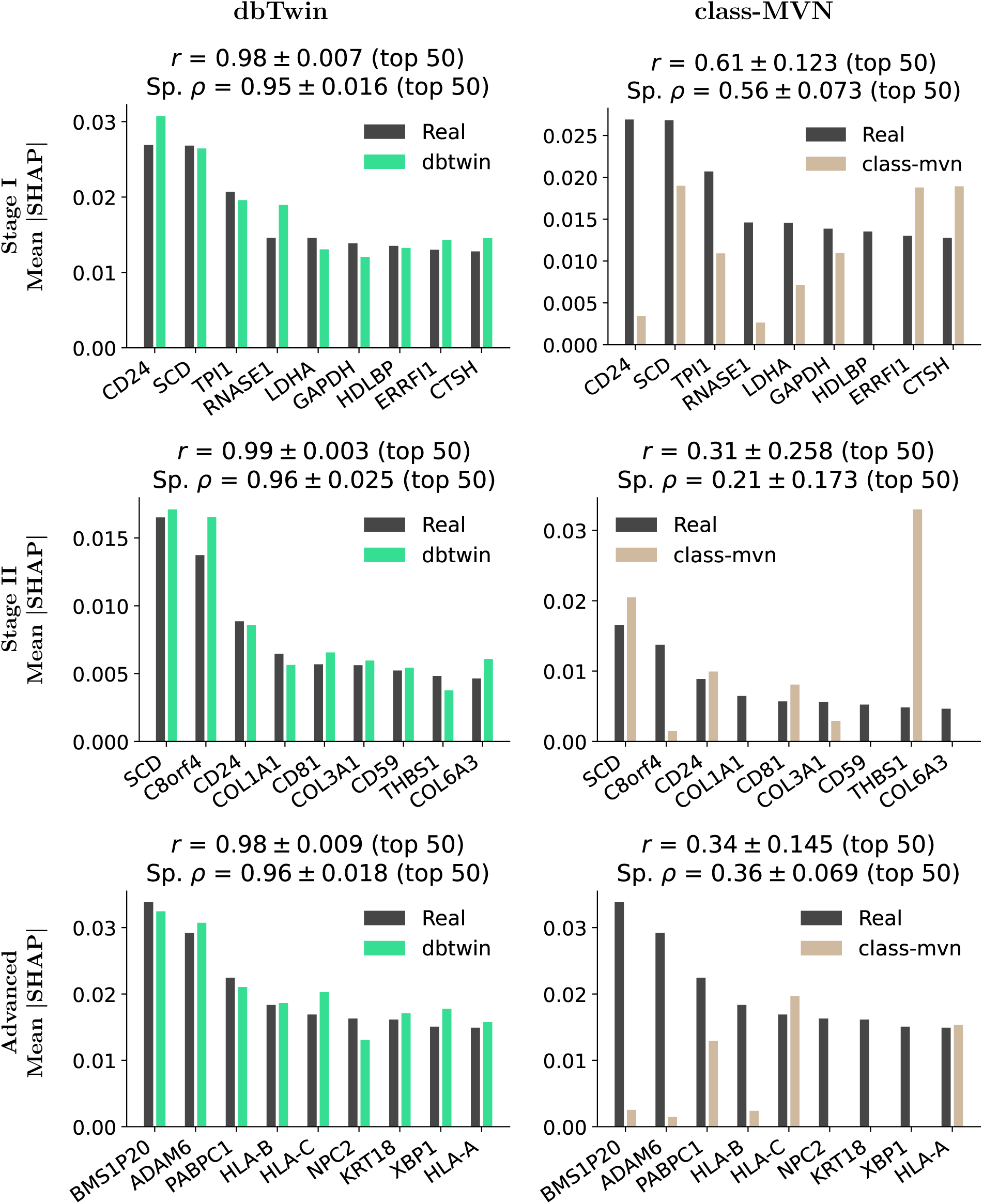
Multiclass ML on LUAD-Stage: SHAP gene-feature concordance: Mean absolute gene-level SHAP weights from classifiers trained on real training (dark) versus synthetic data (colored), both evaluated on the held-out real test set. Annotations indicate Pearson correlation (*r*) for the top 50 genes and Spearman rank correlation (Sp. *ρ*) for the top 50 genes (mean ± SD) over 5 folds. Only median-auc-fold is shown for both models: `dbTwin` (left) and `class-MVN` (right). And, classes: Stage I (top), Stage II (middle), and Advanced (bottom). Complete results for all folds are shown in SI.

### Multivariate Fidelity: Co-expression and UMAP manifold in synthetic cohorts

We focused on whether generative models preserve the high-dimensional correlation structure and global manifold of clinico-transcriptomic data. Here we present results for the `LUAD-EGFR` dataset. Please see the SI for remaining cohorts.

All three generators achieved high Spearman correlations between their synthetic and real coexpression matrices (Fig. 2, left column): `dbTwin` achieved *ρ* = 0.95, `class-MVN` achieved *ρ* = 0.98, and `PCA-CTGAN` achieved *ρ* = 0.92. Results followed a similar trend for all other cohorts with `dbTwin` scoring *ρ*= 0.8–0.95, `class-MVN` scoring *ρ*= 0.98–0.99 and `PCA-CTGAN` scoring *ρ*=0.92– 0.98 - please see SI for details. To understand preservation of low-dimensional structure, we used UMAP [52, 53] to generate 2-dimensional manifolds and maximum-mean discrepancy (MMD) to quantify distributional differences between real (training data) and synthetic manifolds. MMD differentiates generators far more sharply than co-expression correlations: `dbTwin` achieved the lowest MMD (= 0.025), with synthetic samples interleaving tightly with their real counterparts. `class-MVN` maintained acceptable alignment (MMD = 0.19) but showed a noticeable collapse of distributional breadth, with synthetic samples clustering more tightly than real data. `PCA-CTGAN` exhibited severe distributional drift (MMD = 0.53): the synthetic manifold occupies a distinct region of UMAP space. Collectively, all models exhibit preserved co-expression structure for the top variable genes. `dbTwin` provides a slightly better balance across both metrics, preserving both local gene–gene interactions and the low-dimensional distribution simultaneously.

### Differential expression fidelity on binary classification tasks

We assessed DE fidelity between real and synthetic cohorts across the two binary classification cohorts- Sepsis (blood case–control) and `LUAD-EGFR` (EGFR mutation vs wild-type). Quantitative DE gene overlap under moderate stringency (*p*_adj_ *<* 0.05, | log_2_ FC| ≥ 1, baseMean ≥ 10) reveals a clear hierarchy (Fig. 3, Row 1). `dbTwin` recovered 78.7% and 91.8% of real DE genes in `Sepsis` and `LUAD-EGFR` respectively, with a total gene count nearly identical to real data, indicating that the generator neither inflates nor deflates the biological signal. `class-MVN` substantially overestimated DE gene counts (approximately 3× the real count in `Sepsis`), recovering only 64.8% and 43.1% of real genes—the excess corresponds to synthetic artefactual DE signal absent from the real data.

Volcano plot inspection confirms that `dbTwin` synthetic data also preserves the quantitative statistics of DE (Fig. 3, Rows 2-3). We show median performing folds for each model in the volcano plot. All reported Pearson’s correlations are computed over overlapping moderate genes and averaged over 5 folds - see “Methods” for details. Pearson correlations of log_2_FC were near-perfect in both cohorts (`Sepsis`: *r* = 0.99±0.00; `LUAD-EGFR`: *r*= 1.00±0.00), and correlations of − log_10_(*p*_adj_) were high (0.88 ± 0.02 and 0.92 ± 0.05). The synthetic volcano plots are visually indistinguishable from real data: the characteristic asymmetric distribution of up- and down-regulated genes, the cluster of highly significant genes at extreme significance values, and the overall cloud of subthreshold genes are all faithfully reproduced. The slightly lower overlap in `Sepsis` (78.7% vs 91.8% in `LUAD-EGFR`) likely reflects noise near the decision boundary, where log_2_FC and adjusted p-values of borderline genes shift across the threshold under synthetic sampling; nevertheless, the relatively high correlation of both log_2_FC and p-values confirms that `dbTwin` captures the underlying structure, and relaxing the DE thresholds for synthetic data may recover additional boundary genes. Interestingly, contrasting rows 2 and 3 in Fig. 3 shows that the genes in the volcano plot for `dbTwin` are relatively closer together - leading to a smaller angle compared to real data. This is likely because high-fidelity synthetic data has a well-known smoothing effect.

class-MVN captures reasonable fold-change concordance (*r* = 0.96 in `Sepsis`; 0.89 in `LUAD-EGFR`). However, it substantially degrades significance concordance (*r* = 0.68 and 0.24 respectively for − log_10_(*p*_adj_)), consistent with a model that captures broad directional trends but inflates effect sizes and loses the fine-grained p-value architecture. `PCA-CTGAN` showed near-complete signal degradation: fold-change correlations were 0.56 ± 0.26 and 0.19 ± 0.19, and significance correlations collapsed to 0.09 and 0.04 for `Sepsis` and `LUAD-EGFR`, respectively.

**Note:** Given PCA-CTGAN’s relatively poor performance across all DE and UMAP axes, the remainder of the manuscript focuses on `dbTwin` and `class-MVN` and only summary ML results are shown for `PCA-CTGAN` in Fig. 6.

### Machine learning utility on binary classification tasks

We evaluated the utility of synthetic cohorts for predictive modeling using the TSTR (Train-on-Synthetic-Test-on-Real) protocol: classifiers trained on synthetic data are evaluated on held-out real test sets, with classifiers trained on real data (Train-on-Real-Test-on-Real, TRTR) serving as the upper bound. In this subsection, we focus on the two binary tasks; predicting blood sepsis – control from whole-blood RNA-seq data (`Sepsis`) and EGFR mutation status in TCGA-LUAD dataset (`LUAD-EGFR`). For interpretability, we used Logistic regression models with elastic net penalty for all classification tasks. Classifier hyperparameters were selected using cross-validation on the same five-fold stratified split for each real cohort. Importantly, we found that unrestrained optimization of AUC led to over-fitting, resulting in a large number of active gene features with non-zero coefficients. To ensure that downstream SHAP analysis focuses on a compact, interpretable gene set, we constrained hyperparameter selection using a sparsity window. We used the same hyperparameters to train all four models (three synthetic cohorts + real train-set) and tested on the hold-out real test set. Please see Fig. M1 and Section “Methods” for more details.

**Figure M1.**
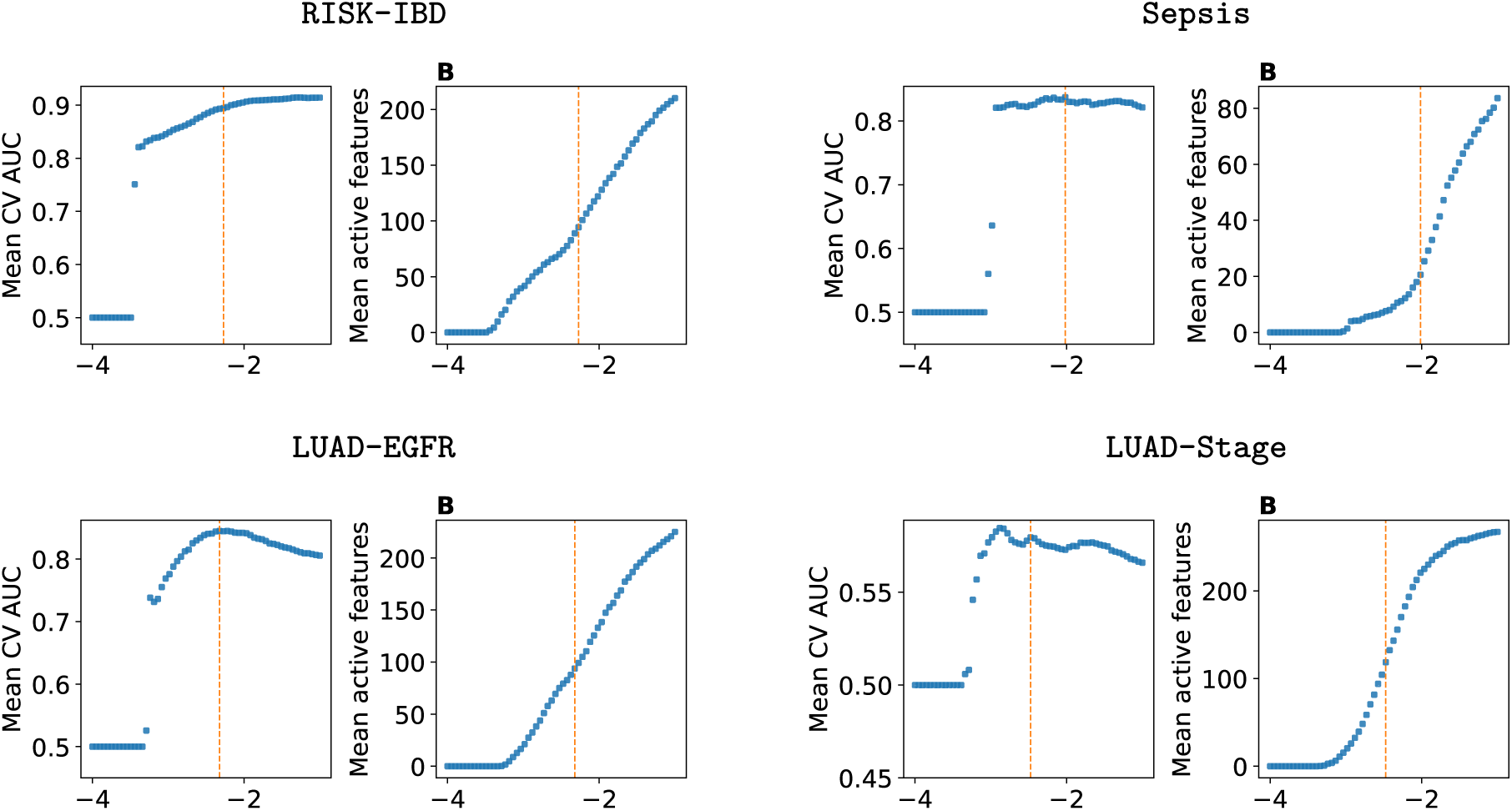
Elastic net hyperparameter selection on real cohorts. Mean 5-fold cross-validation AUC and mean number of active features as a function of log(*C*) for all 4 tasks. The selected *C* (orange dashed line) maximizes mean AUC within the sparsity window 40% *n*_samples_ to 15% *n*_samples_ for `Sepsis, RISK-IBD, LUAD-EGFR`, and `LUAD-Stage`. All cross-validation was done on real cohorts.

Both `dbTwin` and `class-MVN` demonstrated preservation of analytical utility, yielding TSTR AUC scores nearly identical to TRTR in both cohorts (`Sepsis`: TRTR ≈ TSTR ≈0.84; `LUAD-EGFR`: TRTR ≈0.81 versus TSTR ≈0.82; Fig. 5, Top Row) with `class-MVN` exhibiting somewhat higher variance across replicates. Matching AUC alone does not guarantee that a synthetic-trained model has learned the correct biology; a classifier could achieve high discriminative performance by exploiting distributional artefacts in the synthetic data rather than the true gene-level drivers.

We therefore assessed whether synthetic-trained and real-trained classifiers assign importance to the same genes by computing SHAP feature-importance concordance on held-out test data (Fig. 5, Rows 2–3). SHAP analysis is a fairly stringent, model-agnostic approach to quantify contributions of gene-features (*i.e.*, predictors) to model predictive performance on hold-out test data. Measuring concordance between real and synthetic cohort SHAP values is a practical and grounded test of how generalizable synthetic cohorts are: do ML models trained on synthetic data correctly assign similar importance to gene-features when tested on a hold-out real set?

We quantified SHAP concordance using mean absolute SHAP weights correlations computed for each ML model over the top-50 genes on hold-out real cohorts. `dbTwin` achieved high SHAP concordance in `Sepsis` (top-50 genes Pearson’s *r* = 0.90 ± 0.065; Spearman’s *ρ* = 0.85 ± 0.10) recapitulating biologically established sepsis-associated genes (e.g., *CD177*, *S100A9*, *IRAK3*, *SERPINA1*) in the same order as real-trained models, consistent with their well-documented roles in neutrophil activation, innate immune amplification, endotoxin tolerance, and the acute-phase response [54–57]. Similarly, SHAP concordance remained high in `LUAD-EGFR` (top-50 *r* = 0.85±0.036; *ρ* = 0.77 ± 0.08), recovering genes including *ENO1*, *SFTPB*, *KRT7*, and *YWHAZ* in correct rank order, reflecting established roles in glycolytic reprogramming, type II alveolar pneumocyte lineage, adenocarcinoma epithelial identity, and EGFR/PI3K downstream signalling [47, 58, 59].

Importantly, since correlations are averaged across folds, high mean correlation with low standard deviation across cohorts, implying that `dbTwin` accurately preserves fold-wise variance in top predictive genes as well. See SI for SHAP plots on the remaining folds. `class-MVN` achieved moderate concordance in `Sepsis` (*r* = 0.79 ± 0.033; *ρ* = 0.74 ± 0.046) comparable to its performance in other tasks, but degraded substantially in `LUAD-EGFR` (*r* = 0.57 ± 0.131; *ρ* = 0.39 ± 0.067), with the high variance suggesting unstable feature selection across replicates. Our hypothesis for this substantial drop in performance is that `class-MVN` struggles when target classes are highly unbalanced. Results in Figures 7 and 8 add more credence to this view. These results demonstrate that `dbTwin` preserves predictive accuracy and recovers the specific gene-level drivers of classification, a property critical for downstream biomarker discovery.

**Figure 8.**
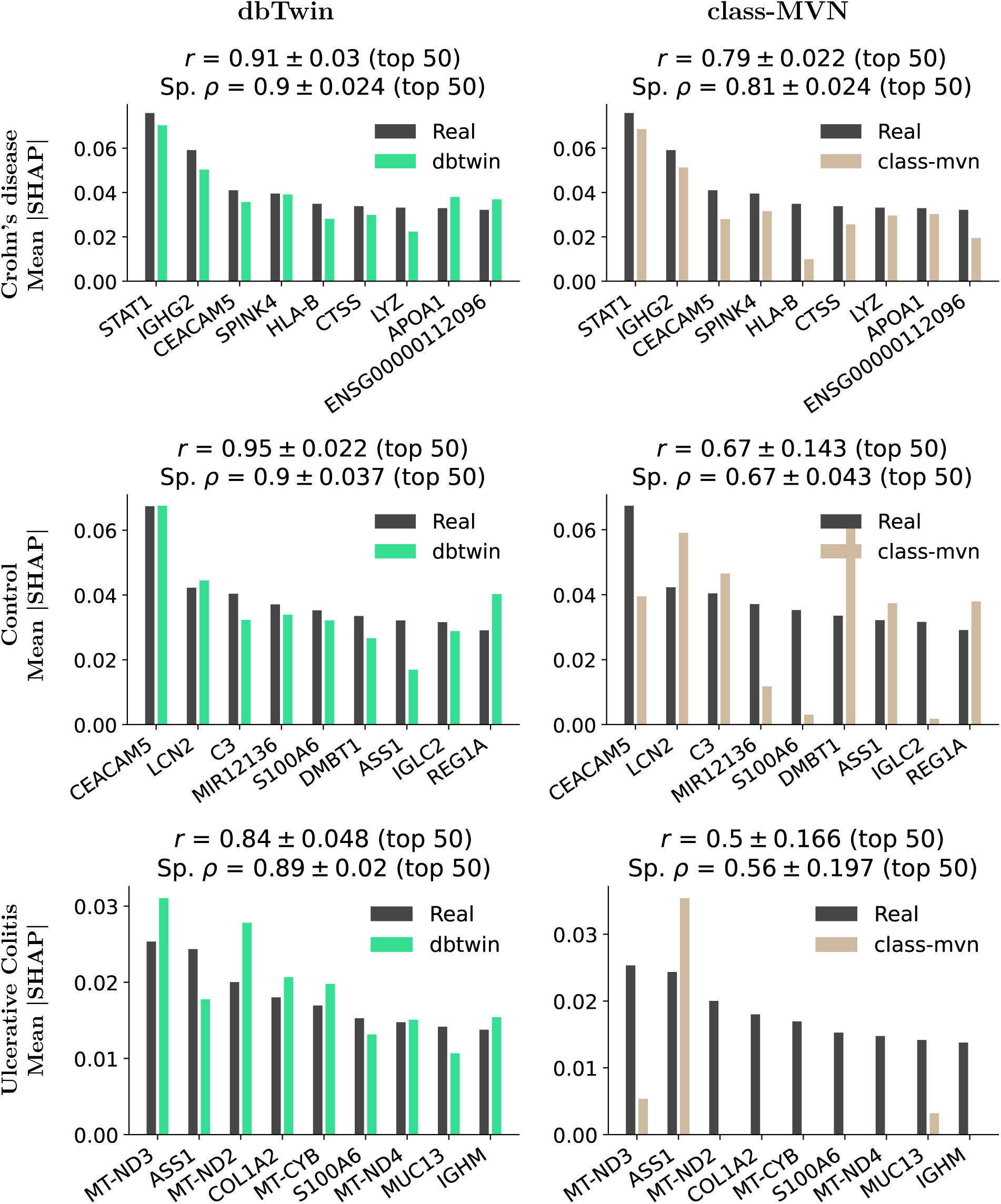
Multiclass ML on RISK-IBD: SHAP gene-feature concordance: Mean absolute gene-level SHAP weights from classifiers trained on real training (dark) versus synthetic data (colored), both evaluated on the held-out real test set. Annotations indicate Pearson correlation (*r*) for the top 50 genes and Spearman rank correlation (Sp. *ρ*) for the top 50 genes (mean ± SD) over 5 folds. Only median-auc-fold is shown for both models: `dbTwin` (left) and `class-MVN` (right). And, classes: Crohn’s disease (top), Control (middle), and Ulcerative colitis (bottom). Complete results for all folds are shown in SI.

### Machine learning utility on multiclass classification tasks

We extended the ML evaluation to two harder multiclass tasks: tumor stage prediction in `LUAD-Stage` (Stage I / Stage II / Advanced) and IBD subtype classification in the `RISK-IBD` pediatric cohort (Crohn’s disease / Ulcerative colitis / Control). We used the same ML model and hyperparameter selection protocol described above with one-vs-rest AUC instead of binary. See “Methods” and Fig. M1 for details.

Both `dbTwin` and `class-MVN` achieved TSTR macro-AUC and macro per-class recall indistinguishable from the TRTR upper bound across both multiclass tasks (Fig. 6), with `class-MVN` showing marginally higher mean AUC on `LUAD-Stage` (∼0.59 vs. ∼0.56) but wider variance across folds. `PCA-CTGAN` substantially under-performed on both tasks.

Since class-averaged SHAP values would be dominated by the majority class, we report SHAP concordance for each class separately. We quantified SHAP concordance using mean absolute SHAP weights correlations computed for each ML model over the top-50 genes on hold-out real cohorts. The tumor-stage classification task on `LUAD-Stage` requires the model to learn separate sets of discriminative genes for each stage (Fig. 7) with the classes in rows 1-3 arranged in descending order of frequency, *i.e.*, Stage I, II and Advanced are the majority, median and minority classes, respectively. `dbTwin` maintained high concordance for all three stages (Stage I: *r* = 0.98 ± 0.007, *ρ* = 0.95 ± 0.016; Stage II: *r* = 0.99 ± 0.003, *ρ* = 0.96 ± 0.025; Advanced: *r* = 0.98 ± 0.009, *ρ* = 0.96 ± 0.018), correctly recovering stage-specific biology. For Stage I, the top drivers were: *CD24*, *LDHA*, and *ERRFI1* (immune evasion, glycolytic reprogramming, and EGFR feedback regulation [60, 61]). For Stage II, *COL1A1*, *COL3A1*, and *THBS1* (desmoplastic remodelling and angiogenic suppression [62]) emerged as dominant features. For Advanced stage, *HLA-A*, *HLA-B*, and *XBP1* were the principal discriminators with downregulation of MHC class I molecules (*HLA-A*, *HLA-B*) denoting canonical immune-evasion mechanism in advanced NSCLC [63]. `class-MVN` showed moderate concordance for Stage I (*r* = 0.61 ± 0.123; *ρ* = 0.56 ± 0.073) but a dramatic drop for Stage II (*r* = 0.31 ± 0.258; *ρ* = 0.21 ± 0.173) and Advanced (*r* = 0.34 ± 0.145; *ρ* = 0.36 ± 0.069), confirming that while `class-MVN` can recover real SHAP weights for some classes, it fails to capture finer-grained biological structure across minority classes.

For IBD subtype prediction, `dbTwin` preserves the correct biological drivers for each subtype as shown in Fig. 8 with the classes in rows 1–3 arranged in descending order of frequency, *i.e.*, Crohn’s disease, Control and Ulcerative Colitis are the majority, median and minority classes, respectively. For Crohn’s disease, `dbTwin` achieved SHAP top-50 *r* = 0.91 ± 0.03 and *ρ* = 0.90 ± 0.024, correctly ranking established IBD genes including *STAT1* (interferon-*γ*/JAK–STAT signalling [64]), *LYZ* (Paneth cell lysozyme [50]) and *SPINK4* (goblet-cell-specific serine protease inhibitor [50]) in near-identical order to the real-trained model. Concordance remained high for the Control class (*r* = 0.95 ± 0.022; *ρ* = 0.90 ± 0.037), with top drivers including *REG1A* (regenerating islet-derived protein) and *DMBT1* (mucosal defence lectin strongly expressed in intact intestinal epithelium), consistent with the stable homeostatic expression defining non-IBD mucosa. Concordance was high for Ulcerative Colitis (*r* = 0.84 ± 0.048; *ρ* = 0.89 ± 0.02), recovering key mitochondrial and barrier features: *MT-CYB* and *MT-ND2* (mitochondrially-encoded respiratory chain subunits, reflecting the epithelial mitochondrial dysfunction characteristic of active UC [65]) and *MUC13* (transmembrane mucin critical for intestinal barrier integrity and dysregulated in UC). `class-MVN` retained moderate concordance for Crohn’s disease (*r* = 0.79 ± 0.022; *ρ* = 0.81 ± 0.024) but degraded for Control (*r* = 0.67 ± 0.143; *ρ* = 0.67 ± 0.043) and Ulcerative Colitis (*r* = 0.50 ± 0.16; *ρ* = 0.56 ± 0.197), with high variance across replicates exhibiting the same decline for median and minority classes noted in LUAD-Stage (Fig. 7).

### Privacy and re-identification risk assessment

Biological fidelity and ML utility are necessary but not sufficient criteria; synthetic cohorts must also demonstrate that they cannot be trivially traced back to individual training subjects as evinced by a growing body of literature pointing to the ease of re-identification using RNA-seq processed counts via reverse-eQTL attacks [13, 17].

We assessed privacy risk using Distance to Closest Record (DCR) on log-transformed and gene-standardized counts, since distances computed in raw-counts space will be dominated by highly expressed genes [66]. While membership inference attacks (MIA) have become a common auditing tool for synthetic tabular health data [67], their threat model assumes an adversary who already possesses a target individual’s real record and seeks to confirm its inclusion in the training set. For non-identifiable gene expression data governed under the Common Rule, this assumption is largely unrealistic, rendering the threat model circular and membership confirmation moot. We further note that differential privacy, often proposed as a formal guarantee for synthetic data release, faces well-documented practical limitations: meaningful privacy budgets (*ɛ* ≤ 1) destroy utility in high-dimensional settings, while the large *ɛ* values required to preserve utility offer negligible protection [68, 69]. DCR-based metrics, by contrast, directly measure whether synthetic records are implausibly close to real training samples - a more interpretable and threat-model-agnostic measure of re-identification risk.

We evaluate re-identification risk using Distance to Closest Record (DCR) distributions [70]. We summarize these distance distributions using a p5 ratio, defined as the 5th percentile of synthetic-to-training distances divided by the 5th percentile of synthetic-to-holdout distances, with values near 1 indicate no preferential proximity to training records. For both `dbTwin` and `class-MVN`, DCR distributions show substantial overlap between syn → train and syn → test across all datasets, confirming the absence of gross memorization (Fig. 9). `class-MVN` achieved near-perfect equivalence (DCR p5 ratio at 1.00), while `dbTwin` showed slightly lower p5 ratios 0.90–0.94, reflecting marginally shorter synthetic-to-training distances consistent with its higher fidelity generation, though synthetic counts remained well separated from training records. Together, these results confirm that neither generator produces memorized copies of real expression profiles.

**Figure 9.**
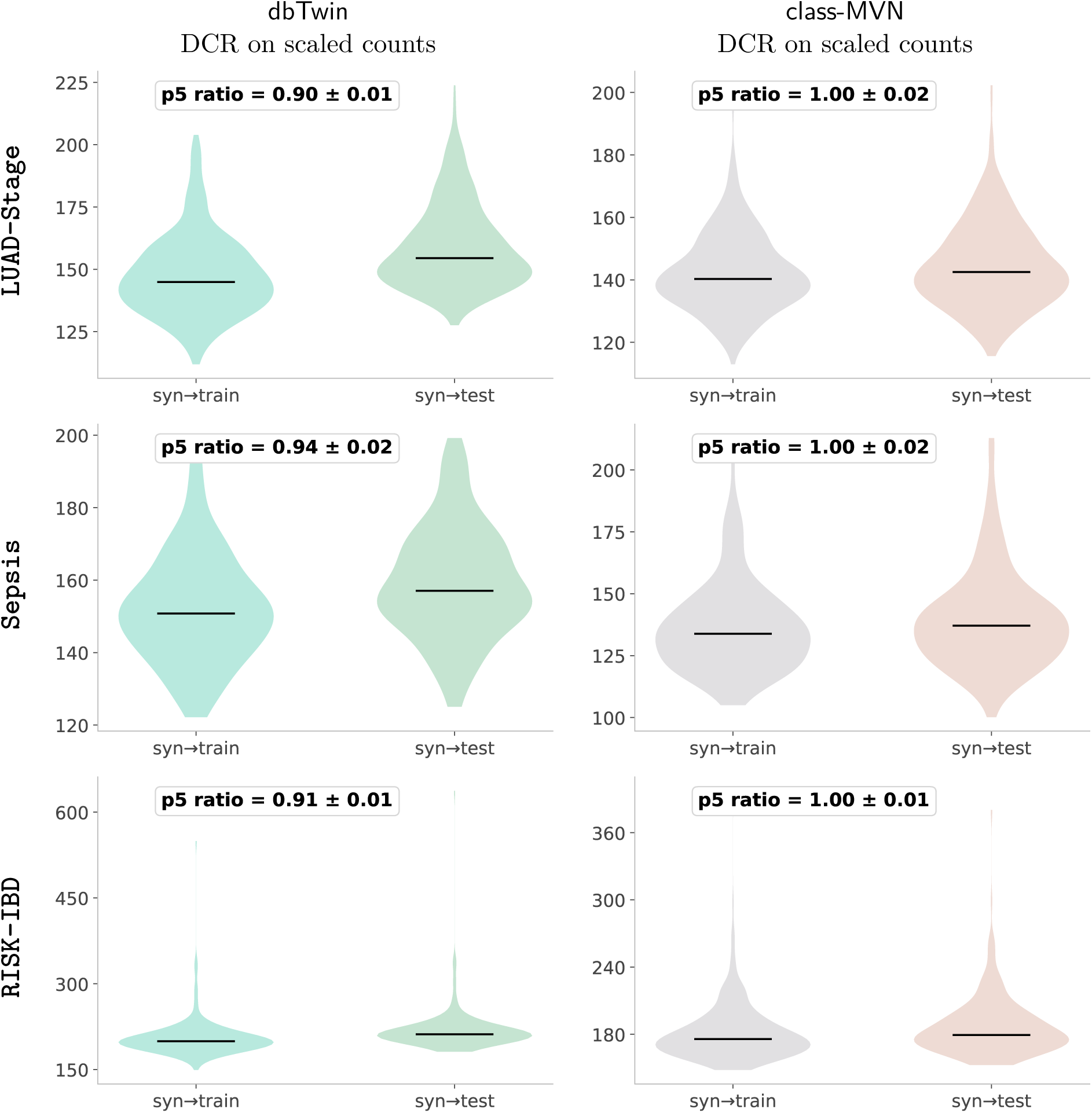
Distance to closest record (DCR) privacy metric. Results shown for three dataset-target pairs (rows) and two generators (columns). `LUAD-EGFR` results are in SI. DCR violins show distance for each synthetic sample to its nearest neighbor in the real training set (syn→train, blue) versus the held-out test set (syn→test, orange). The p5 ratio (annotated per panel) is the 5th percentile of syn→train distances divided by the 5th percentile of syn→test distances; values near 1, with overlapping distance distributions indicate no excess proximity to training data. Both generators pass this check across all three cohorts.

## Discussion

We set out to evaluate synthetic clinico-transcriptomic cohorts against a biologically grounded multi-axis benchmark, and to assess whether single-metric evaluation paradigms are sufficient to detect failures across these axes. The results presented here suggest that both questions can be answered affirmatively, but with important caveats that merit careful discussion.

### A discovery-aware, transcriptome-scale benchmark

The recent comprehensive benchmark by Öztürk et al. represents an important step toward principled evaluation of synthetic RNA-seq generators, comparing 11 models across fidelity and utility metrics including feature importance [46]. However, that benchmark was limited to at most 1,000 genes per cohort, and its utility metrics, while multi-axis, remain at a granularity that may not surface all failure modes relevant to translational workflows. Our work builds on and extends that framework in two specific ways.

First, our benchmark operates at realistic transcriptome scale (approximately 30,000 to 50,000 genes). This distinction matters because translational analyses rarely begin with a pre-filtered gene set, and a generator that performs well on a curated subset may fail when asked to reproduce the joint distribution over the full transcriptome. Second, we deepen the evaluation axes to capture failure modes that are specifically relevant to how clinico-transcriptomic cohorts are used in discovery workflows. For differential expression, we move beyond gene-set overlap to quantify gene-wise concordance of both log2 fold-change magnitude and adjusted p-value significance. This matters because a generator that recovers the correct set of DE genes but inflates their significance (as *`class-MVN`* does, producing approximately 3x the real DE gene counts in the `Sepsis` cohort) will mislead downstream hit-list prioritization. For ML utility, we move beyond aggregate AUC to per-class SHAP feature-attribution concordance computed on held-out real data. This addition demonstrates the need for multi-axis fidelity metrics since a synthetic cohort can easily match realdata AUC while learning biologically incorrect features. `class-MVN` illustrates this clearly in the LUAD-Stage task, where it achieves comparable AUC to real while its per-class SHAP concordance drops to *r* = 0.31–0.34 for Stage II and Advanced classes.

### SHAP concordance as a necessary condition for translational relevance

SHAP concordance deserves particular emphasis because it addresses a failure mode invisible to standard metrics. A synthetic cohort used for differential expression analysis or biomarker discovery in a data-sharing context must produce not only accurate predictions but also the correct set of gene-level drivers. Critically, SHAP values are computed on the hold-out test data not used for synthetic cohort generation; therefore, SHAP concordance goes one-step further and directly measures the generalization of ML models trained on the synthetic data to unseen real-world cohorts.

This generalization held across all four evaluation cohorts. dbTwin’s synthetic-trained classifiers recovered the same top gene-level drivers as real-trained models in both binary tasks (Pearson’s *r* = 0.85–0.90 on top-50 genes) and maintained high per-class concordance in both multiclass settings (*r* = 0.84–0.99 across all `RISK-IBD` and `LUAD-Stage` classes). Strikingly, `dbTwin` exhibited relatively high mean correlation with low std across cohorts values implying that `dbTwin` accurately preserves fold-wise variance in top predictive genes as well; this may have implications for federated synthetic data generation approaches [71]. `class-MVN`, in contrast, sustained moderate SHAP concordance in the `Sepsis` task but degraded sharply in multiclass and minority class settings (*r* = 0.31–0.61 for `LUAD-Stage`; *r* = 0.50–0.79 for `RISK-IBD`) despite closely tracking real-AUC values across cohorts, illustrating precisely the failure mode that single-metric evaluation would miss.

These results suggest that sufficiently high SHAP concordance, if confirmed across datasets and conditions, could support a synthetic-first discovery workflow: a data custodian releases a synthetic version of a controlled-access cohort under open or minimal-access terms; an external researcher uses it for exploratory DE analysis, biomarker discovery, and feature prioritization; candidate gene sets are then taken to a formal data access request for validation on the real cohort. This two-stage approach could reduce the number of access requests that fail to yield results, while giving researchers a more informative starting point than metadata or summary statistics alone.

### Privacy: what DCR captures and what it does not

DCR as an empirical privacy metric is directly interpretable and generator-agnostic. Under this metric, both `dbTwin` and `class-MVN` pass the empirical test across all three datasets, with `dbTwin` showing slightly lower p5 ratios (0.90–0.94) while `class-MVN` achieved near-perfect equivalence (1.00 ± 0.01 to 1.00 ± 0.02). We explicitly note that the DCR analyses establish that synthetic samples are not memorized copies of real expression counts.

It is important to test this because synthetic data can score exceedingly well on utility metrics by merely memorizing real data. However, as a caveat, DCR-based analyses are best understood as a necessary but not sufficient condition for privacy. In practice, a truly private synthetic dataset needs to demonstrate low re-identification risk in the context of an appropriate threat model. A comprehensive, model-agnostic privacy analysis under realistic adversarial threat models is the subject of a forthcoming companion study.

### Ultra-high dimensionality fundamentally challenges deep generative models

An empirical finding across all evaluation axes is that a non-deep-learning generator, `dbTwin`, and a statistical model, `class-MVN`, consistently outperformed a GAN-based generator operating in PCA-compressed space - PCA-CTGAN. This outcome is informative about the structure of the problem itself;RNA-seq count matrices comprise 20,000–50,000 genes with complex covariance structure driven by co-expression modules, pathway organization, and tissue-specific expression programs [45, 72] and provide additional evidence [46, 73, 74] that deep-learning-based generative models may be ill-suited for large-scale biomedical datasets.

### Limitations

Several additional limitations should be noted. First, our ML evaluation uses a single classifier family (elastic-net logistic regression); while this choice was motivated by interpretability and ensures that SHAP values reflect well-understood gene-level weights that capture synthetic-trained model generalization to unseen real data, whether synthetic cohorts exhibit similar performance on tree-based or deep-learning classifiers remains an open question. Second, the current evaluation axes all require a single target column for DE or ML classification, which may be insufficient for translational workflows that occasionally require accurate joint modelling of continuous and categorical clinical covariates alongside expression data. Third, we limited our analyses to DE and ML, future work should examine whether synthetic cohorts can capture the pathway-level structure of real cohorts. Fourth, all datasets examined here are from single-platform, relatively homogeneous processing pipelines. Real-world RNA-seq data frequently contains batch effects from different sequencing platforms, library preparation methods, and processing centers. Harmonizing multisite federated datasets via synthetic data to enable multi-party collaboration will be an important challenge to tackle. Lastly, all datasets examined here are bulk RNA-seq with relatively clean clinical annotations; extension to pseudo-bulk and single-cell RNA-seq, where the dimensionality and sparsity challenges are even more acute, will be an important next step.

### Practical guidance for data custodians

Synthetic cohorts are likely sufficient for exploratory analysis, hypothesis generation, method development, and educational use, where approximate fidelity is adequate. Real data remains necessary for regulatory submissions, clinical validation, and rare-variant analyses, where tail behavior and exact record-level statistics matter. Before data custodians release synthetic cohorts, they should be validated on both biological fidelity and empirical privacy risk using a multi-axis framework such as the one presented here; the benchmark code is released to support this directly.

### Future directions

Looking forward, this benchmark provides a foundation that can be extended along several axes. The most pressing is a rigorous privacy evaluation grounded in the specific attack models for gene-expression datasets. Beyond privacy, the framework should be extended to evaluate synthetic data utility for survival analysis, pathway enrichment, and network inference. A third interesting direction is evaluating `dbTwin` and other models’ performance on pseudo-bulk data from scRNA-seq datasets - noting that pseudo-bulk datasets share schema and dimensionality with bulk RNA-seq cohorts.

## Author Contributions

AN: Conceptualization, methodology, software implementation, data acquisition and pre-processing, development and implementation of generative models, benchmarking and evaluation protocols, literature review, and principal manuscript drafting. SS: Interpretation and biological relevance of the results; contributed to the Results and Discussion sections of the manuscript.

## Acknowledgments

AN thanks Ashish Patel for his support and help. We thank the recount3 team for providing accessible RNA-seq data resources. Computational resources were supported by Microsoft Azure Cloud startup credits.

## Code and data availability statement

All benchmark code and summary statistics for reproducing figures are available at https://github.com/Nanda-Aditya/rna-syn-bench [75]. All real and generated synthetic cohorts are released at Synapse:syn75080394.

## Methods

### Datasets and Use-Cases

We evaluated synthetic data generators across four clinico-transcriptomic cohorts spanning oncology, sepsis and pediatric IBD conditions (Table 1). To probe a diverse set of biological questions, we considered two binary classification problems and two multiclass classification problems. DE analysis was only conducted on the two binary tasks (`LUAD-EGFR` and `Sepsis`) and predictive ML tasks on all 4 cohorts.

For each dataset-target pair, we generated five independent random train-test splits using stratified sampling to ensure class-proportion balance across folds. This stratification was applied to both binary and multiclass (3-class) targets. All generative models were trained exclusively on the training portion, with the held-out test set reserved for downstream evaluation. Expression data were stored as raw integer counts to preserve the statistical properties of RNA-seq measurements.

### Generative models

#### dbTwin

dbTwin (dbTwin, Inc.) is a proprietary, non-deep-learning generative engine that extends linear algebra-based synthetic data generation methods [33, 34] to bulk RNA-seq cohorts. It operates directly in count space at full transcriptome scale (∼50 000 genes) and conditions synthesis on per-sample biological targets including both the classification label and paired clinical covariates.

#### class-MVN

We created a modified version of `class-MVN` [46] using a low-rank approximation that natively scales up to transcriptomic scale. The original `class-MVN` works by fitting a simple multivariate normal distribution to preserve gene-gene correlations within each target class (in log space). We extended this method by using a randomized SVD to project all samples into a low-rank linear subspace, fitting a per-class Gaussian in that reduced space, sampling, and projecting back to gene space. The SVD rank was set to *k* = min ⌊0.6 *n*_train_⌋, *n*_min-class_ − 1, where *n*_min-class_ is the number of samples of the smallest class, to avoid rank deficiency within classes. `class-MVN` does not model clinical covariates beyond the target label. The generation code for `class-MVN` (and PCA-CTGAN) is available at https://github.com/Nanda-Aditya/rna-syn-bench.

#### PCA-CTGAN

We created a modified version of CTGAN [35] able to train on and sample from full gene expression counts and paired clinical covariates. Deep-learning architectures like CTGAN have training times that grow quadratically with the number of columns; training on more than ≈ 500 columns therefore becomes extremely slow and costly. To overcome this, we first log-transformed counts (log_2_(*x* +1)) and applied PCA, retaining *k* = max(2, ⌊0.6 *n*_train_⌋)components. Clinical covariates were concatenated with the PCA scores, and the combined matrix was passed to CTGAN (300 training epochs). Synthetic PCA vectors were inverse-transformed, exponentiated, and rounded to recover integer count matrices.

### Differential Expression Fidelity

To assess whether synthetic cohorts preserve biologically meaningful gene expression signals, we performed differential expression (DE) analysis independently on both real and synthetic datasets using DESeq2 [76, 77]. Prior to DE analysis, genes with a median absolute deviation (MAD) below the 20th percentile of the training set were excluded to remove near-constant features that add noise without biological signal. For each dataset, we fit a negative binomial generalized linear model with the clinical variable of interest (EGFR mutation–wild-type for `LUAD-EGFR` and blood sepsis case–control for `Sepsis`) as the design factor, then extracted per-gene log_2_ fold-changes and padj-values.

For brevity, we evaluated concordance between real and synthetic DE results at a moderate threshold (adjusted *p <* 0.05, |log_2_FC| ≥ 1, baseMean ≥ 10). For Figures 3 and 4, we computed the overlap between the real and synthetic DE gene sets as a percentage of the real set averaged over all 5 folds. All volcano plots were plotted using log_2_ fold changes and adjusted p-values (−*log*10(*padj*)). To quantify agreement between real and synthetic cohorts, we calculated the Pearson’s correlation of log_2_ fold-changes and padj on overlapping moderate stringency genes. Higher correlations paired with strong gene-set recapitulation show that the synthetic cohort faithfully recapitulates the direction and magnitude of differential expression.

### Machine Learning Evaluation

#### Hyperparameter Optimization

Classifier hyperparameters were selected entirely on real data to avoid any information leakage from the synthetic generators into the evaluation. We used cross-validation on the same five-fold stratified split for each dataset employed for synthetic-data generation. For each fold, gene-expression values were log-transformed (log_2_(*x* + 1)) and the top *N* most variable genes were selected by median absolute deviation (MAD), computed on the training fold to prevent test-set leakage. The feature count was set to *N* = min 270, 0.9 *n*_train_), ensuring the gene set never exceeds 90% of the training sample count. A logistic regression classifier with elastic-net penalty was then trained using the saga solver (maximum 5 000 iterations). For computational ease, we fixed *ℓ*_1_ ratio at 0.05. The regularization strength *C* was swept over a predefined grid; for each candidate value, the macro-averaged AUC (one-vs-rest) was recorded on the corresponding test fold, together with the number of active gene-features, *i.e.* genes with non-zero coefficients. For binary tasks (`Sepsis, LUAD-EGFR`) the standard binary AUC was used instead.

Importantly, we found that unrestrained optimization of AUC led to over-fitting, resulting in a large number of active gene features with non-zero coefficients. To ensure that downstream SHAP analysis focuses on a compact, interpretable gene set, we constrained hyperparameter selection to a sparsity window. For each dataset, a dataset-specific threshold *n*_thres_ was set to approximately 40% of the training set size: *n*_thres_ = 48 (`Sepsis`), 100 (`RISK-IBD`), and 160 for both TCGA-LUAD tasks (`LUAD-EGFR, LUAD-Stage`). For one cohort, this led to extremely sparse solutions, so we added a lower-bound of 15% of train-set size to all 4 cohorts. The optimal *C* was then chosen as the value yielding the highest mean AUC across the five folds with number of active gene-features within this sparsity window.

#### Train-on-Synthetic, Test-on-Real Evaluation

To quantify the downstream utility of each synthetic dataset, we adopted a *train-on-synthetic, teston-real* (TSTR) protocol. For every combination of generator and cross-validation fold, a logistic regression model (elastic-net, with the *C* selected in the hyperparameter optimization above) was trained on the synthetic training set and evaluated on the corresponding held-out *real* test fold. Gene-expression features were log-transformed (log_2_(*x* + 1)) and filtered to the top genes ranked by MAD, consistent with the preprocessing applied during hyperparameter selection. The same classifier was also trained on the real training fold to provide an upper-bound reference. For each fold we recorded:

1. **Macro-averaged AUC** (one-vs-rest for multiclass tasks; binary AUC for two-class tasks), providing an overall measure of discriminative performance.
2. **Macro-averaged per-class recall**, capturing each generator’s ability to preserve class-specific signal.
3. **SHAP values on hold-out test** capturing predictive performance of models trained on synthetic and real train data

All metrics were averaged over the five folds; we report mean ± standard deviation. To assess whether synthetic data preserve the biological feature importances learned from real samples, we computed SHAP values [78] on the held-out real test data for every trained model. We then measured the concordance between the SHAP-derived feature rankings of the real-trained and each synthetic-trained model using two complementary correlation measures: Pearson’s *r* over the top 50 genes by mean absolute SHAP value, and Spearman’s *ρ* over the top 50 genes. Both correlations were averaged (± standard deviation) across the five folds. High SHAP concordance indicates that a synthetic dataset not only matches overall predictive accuracy but also recovers the same gene-level drivers of classification while preserving fold-wise variance, a property critical for downstream biomarker discovery and translational interpretation.

### Privacy Risk Assessment

#### Expression-Space Privacy Metrics

To assess whether synthetic gene-expression profiles memorize or closely replicate real patient samples, we computed closest-record distances in standardized expression space. All gene-expression columns were log-transformed and *z*-scored using the training-set mean and standard deviation. Both synthetic data and held-out test data were transformed with the same statistics to ensure a common scale.

For each synthetic sample, we measured the Euclidean distance to its nearest neighbors in two reference sets: (i) the train set from which the synthetic cohort was generated and (ii) the held-out test cohort. Because the training and test sets differ in size, we sub-sampled the train-set to match the test-set size in each of *R*=5 independent rounds and averaged the resulting distance vectors, yielding mean distance-to-training *d*_S*→*trn_ and distance-to-test *d*_S*→*tst_ for every synthetic record.

We summarize these distributions with paired violin plots (Figure 9) that overlay *d*_S*→*trn_ and *d*_S*→*tst_ for each generator and dataset combination. A generator that has not memorized training samples should produce comparable distance distributions to both reference sets; systematic shifts toward smaller *d*_S*→*trn_ indicate potential overfitting or privacy leakage. As a scalar summary metric, we report the *5th-percentile distance ratio*:

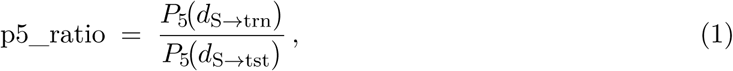

where *P*_5_(·) denotes the 5th percentile. Values near 1 indicate that the closest synthetic-real pairs are equally distant from training and test records, whereas values significantly below 1 signal that some synthetic samples sit closer to training records than would be expected by chance.

### Unsupervised Utility Metrics

Gene-gene correlation structure was assessed using Pearson correlation matrices for the top 300 highest-variance genes. Hierarchical clustering leaf ordering was computed from real data and applied consistently across synthetic matrices to enable visual comparison.

UMAP embeddings were computed jointly on real and synthetic expression data (all generators combined in a single fit) to enable direct coordinate comparison. Genes below the 40th MAD percentile of the training set were excluded prior to embedding, retaining the top 60% most variable genes. Embeddings used log-transformed counts with hyperparameters: `n_neighbors= 15, min_dist= 0.1, metric=euclidean, n_components= 3`; the first two components are visualized. Maximum mean discrepancy (MMD) between real and synthetic manifolds was computed on the 2-dimensional projected coordinates using an RBF kernel (default bandwidth *γ* = 1*/*2):

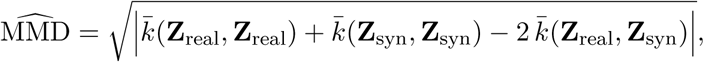

where *k̄*(**A**, **B**) denotes the mean kernel value over all row pairs.

### Software and Implementation

Analyses were implemented in Python using scikit-learn, numpy and scipy for machine learning, pyDESeq2 for differential expression, SHAP for feature attribution, and UMAP for dimensionality reduction. Statistical computations used SciPy and NumPy.

