## Supplementary Information for "Synthetic RNA-seq cohorts for data sharing: a discovery-aware benchmark at transcriptome scale"

### Multivariate Fidelity — TCGA-LUAD EGFR

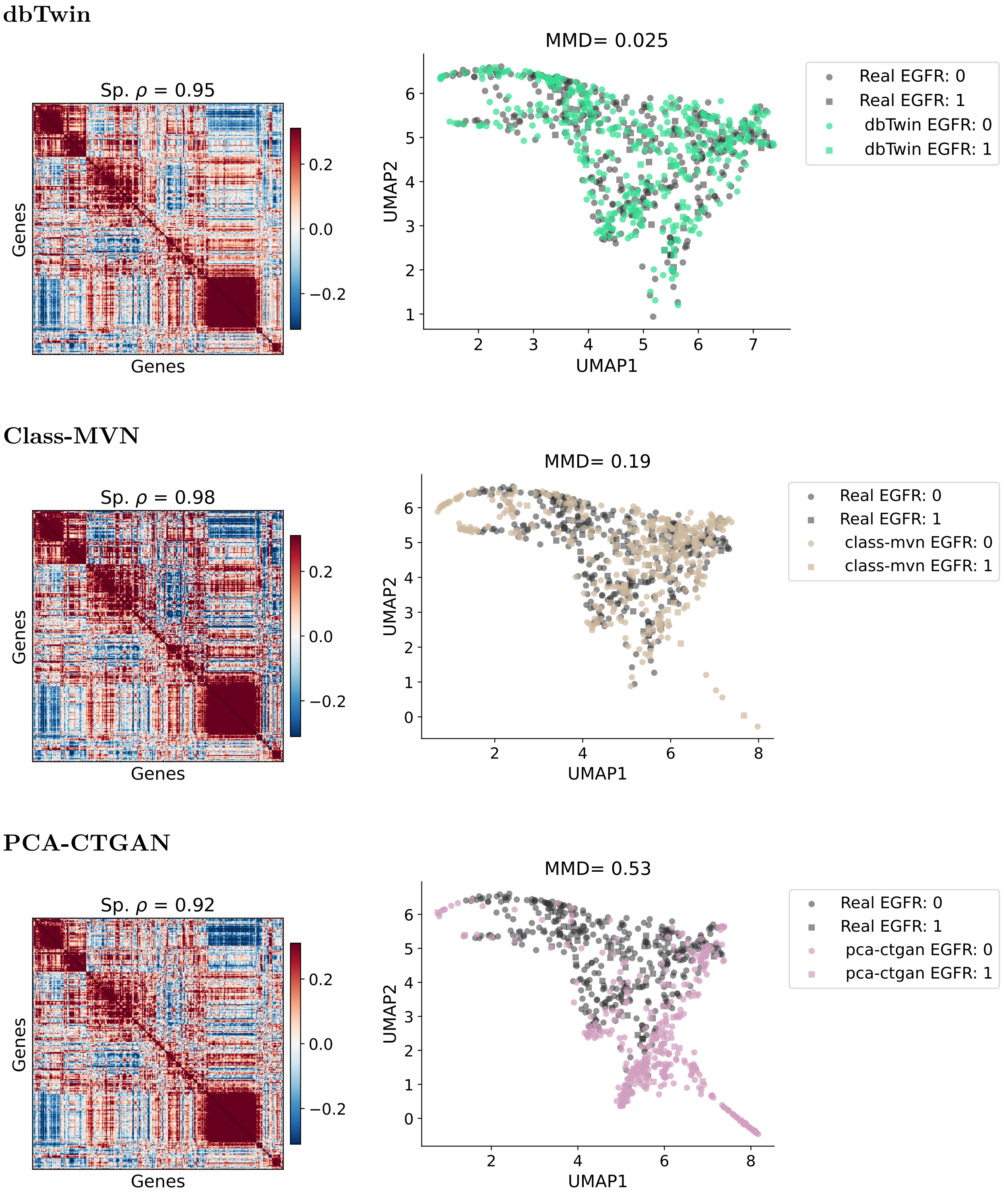


**Figure 1. Multivariate fidelity assessment for the TCGA-LUAD EGFR mutation dataset. (a–c)** Gene co-expression correlation matrices comparing real data (left triangle) versus synthetic data (right triangle) for the top 300 most variable genes. **(d–f)** UMAP embeddings (components 1 vs 2) showing training data (grey points) overlaid with synthetic data (colored points) for each generator.

### Multivariate Fidelity — Sepsis

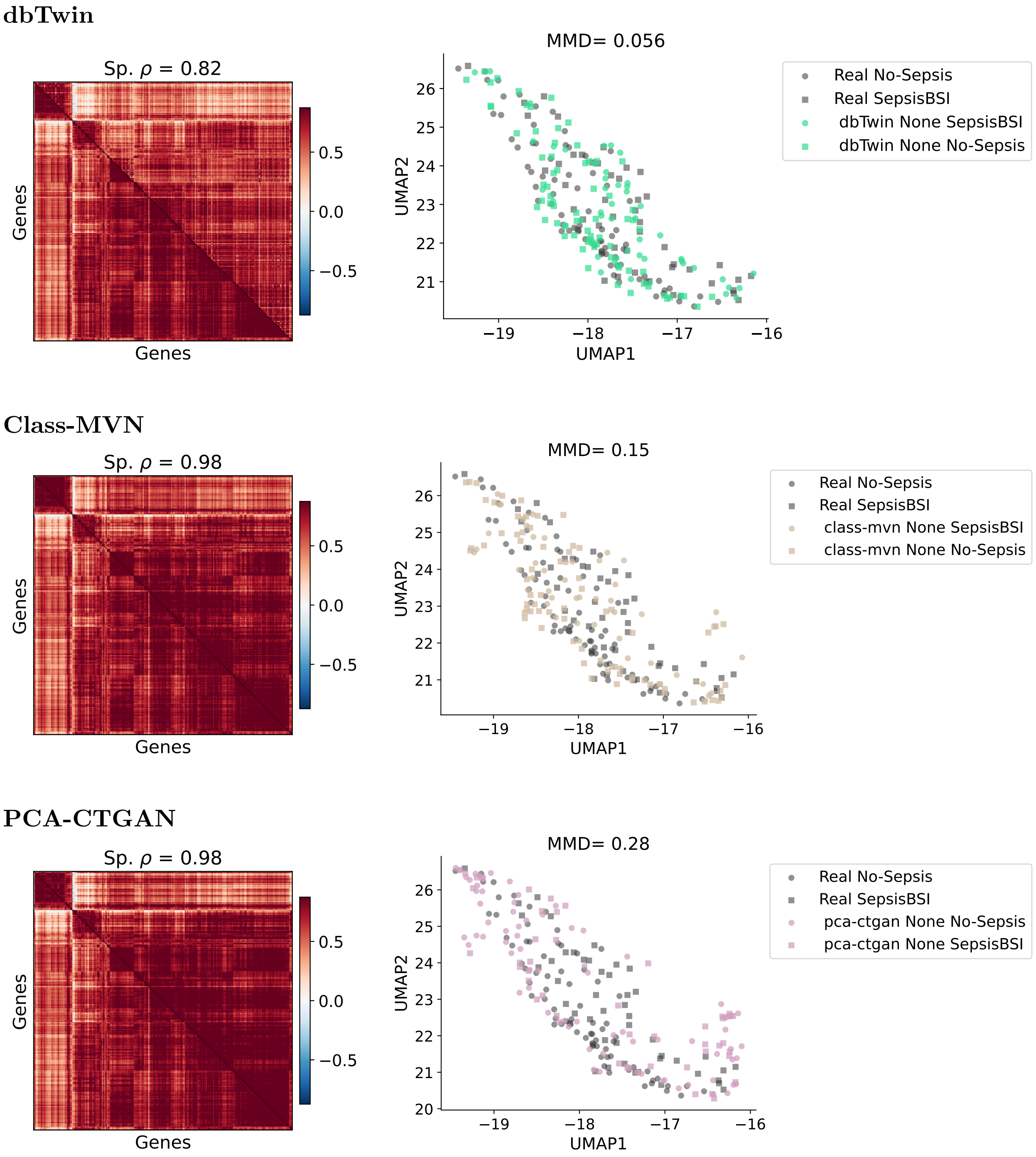


**Figure 2. Multivariate fidelity assessment for the Sepsis dataset. (a–c)** Gene co-expression correlation matrices comparing real data (left triangle) versus synthetic data (right triangle) for the top 300 most variable genes. **(d–f)** UMAP embeddings (components 1 vs 2) showing training data (grey points) overlaid with synthetic data (colored points) for each generator.

### Multivariate Fidelity — RISK

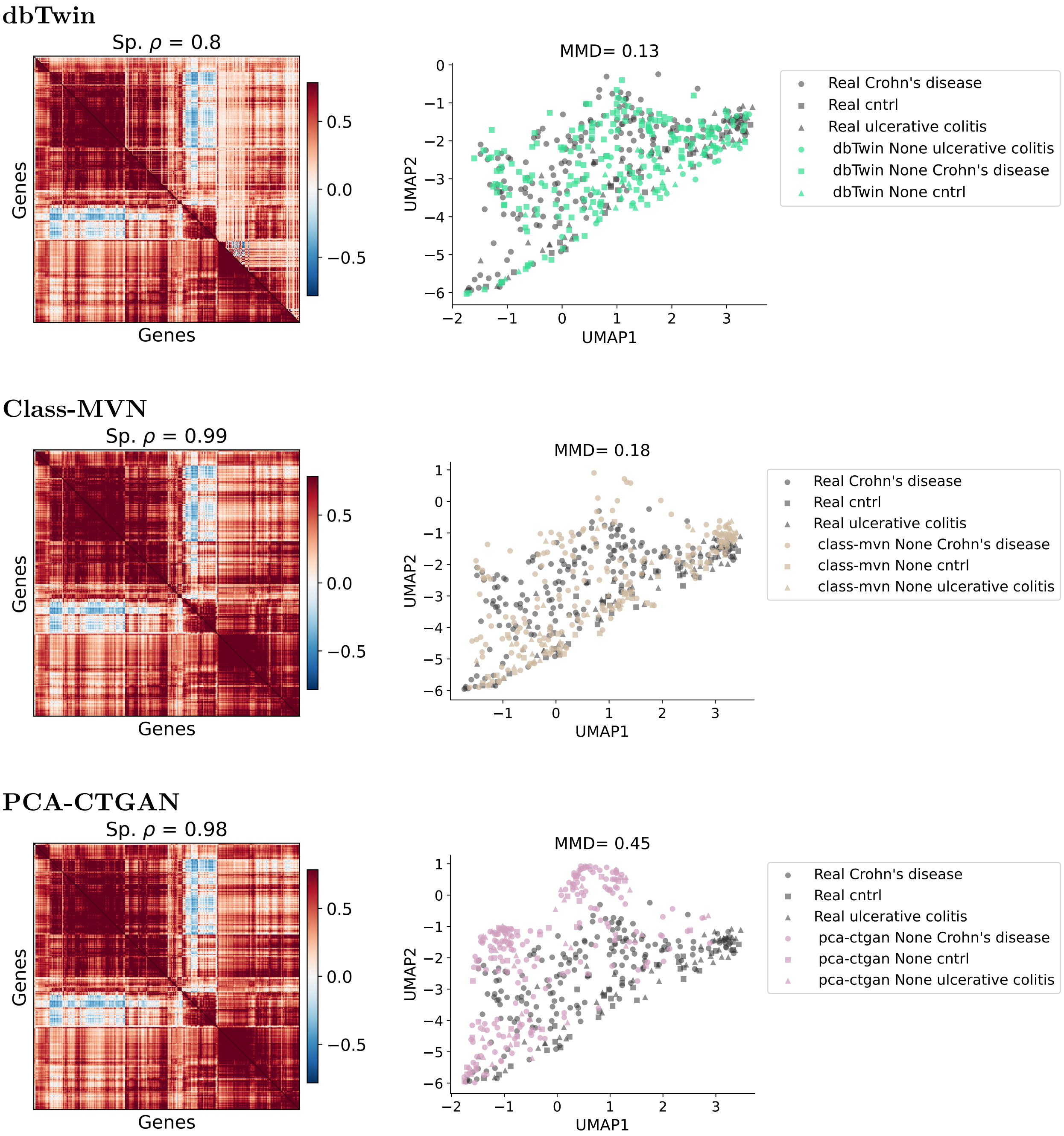


**Figure 3. Multivariate fidelity assessment for the RISK pediatric dataset.** **(a–c)** Gene co-expression correlation matrices comparing real data (left triangle) versus synthetic data (right triangle) for the top 300 most variable genes. **(d–f)** UMAP embeddings (components 1 vs 2) showing training data (grey points) overlaid with synthetic data (colored points) for each generator.

### SHAP Analysis — Sepsis Classification

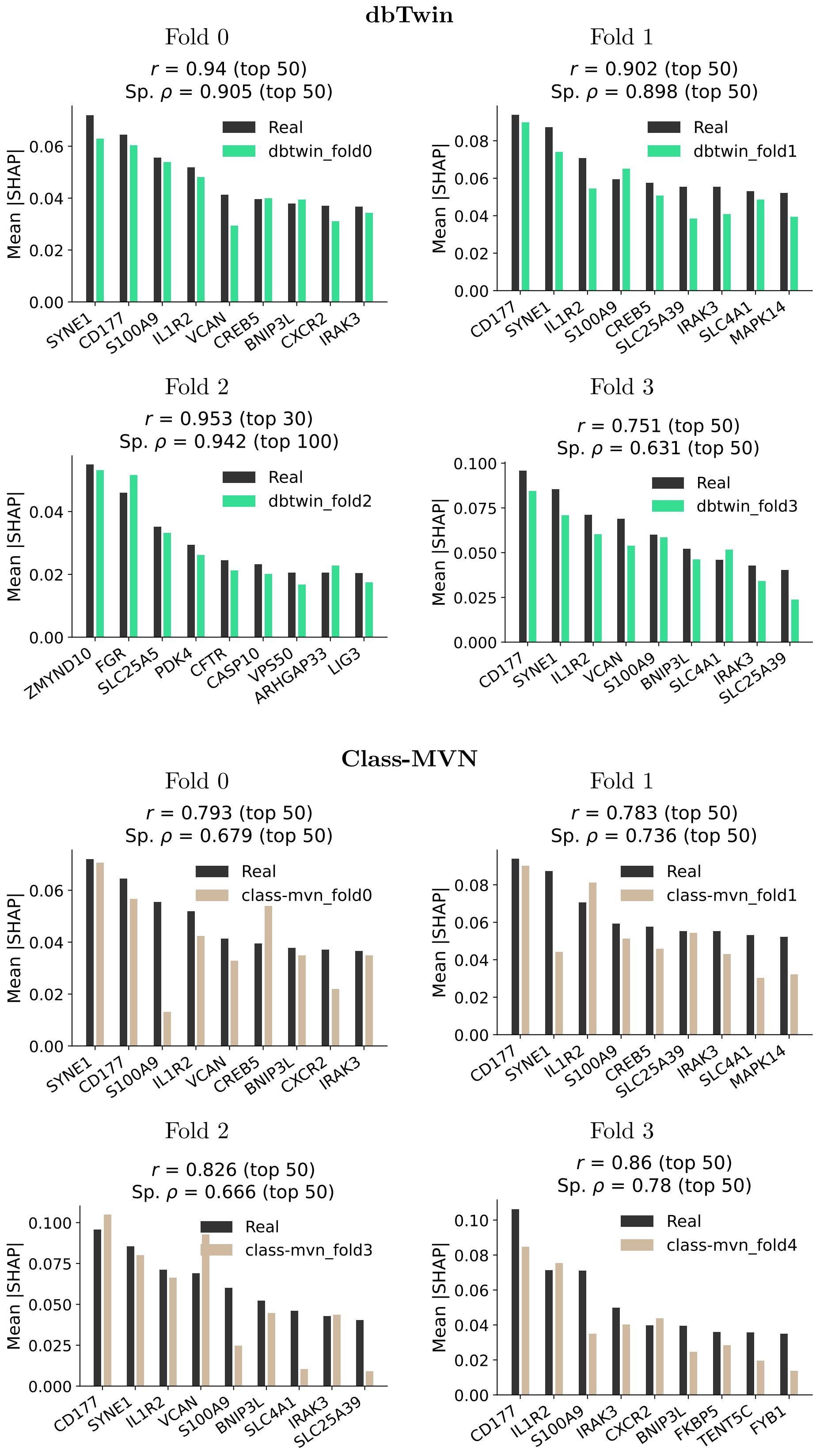


**Figure 4. SHAP gene-feature concordance for Sepsis classification dataset in 4 remaining folds not shown in main text.** Top panel: **dbTwin**. Bottom panel: **Class-MVN**. Each plot shows the top gene-features that contribute most to the classification decision.

### SHAP Analysis — TCGA-LUAD EGFR Mutation

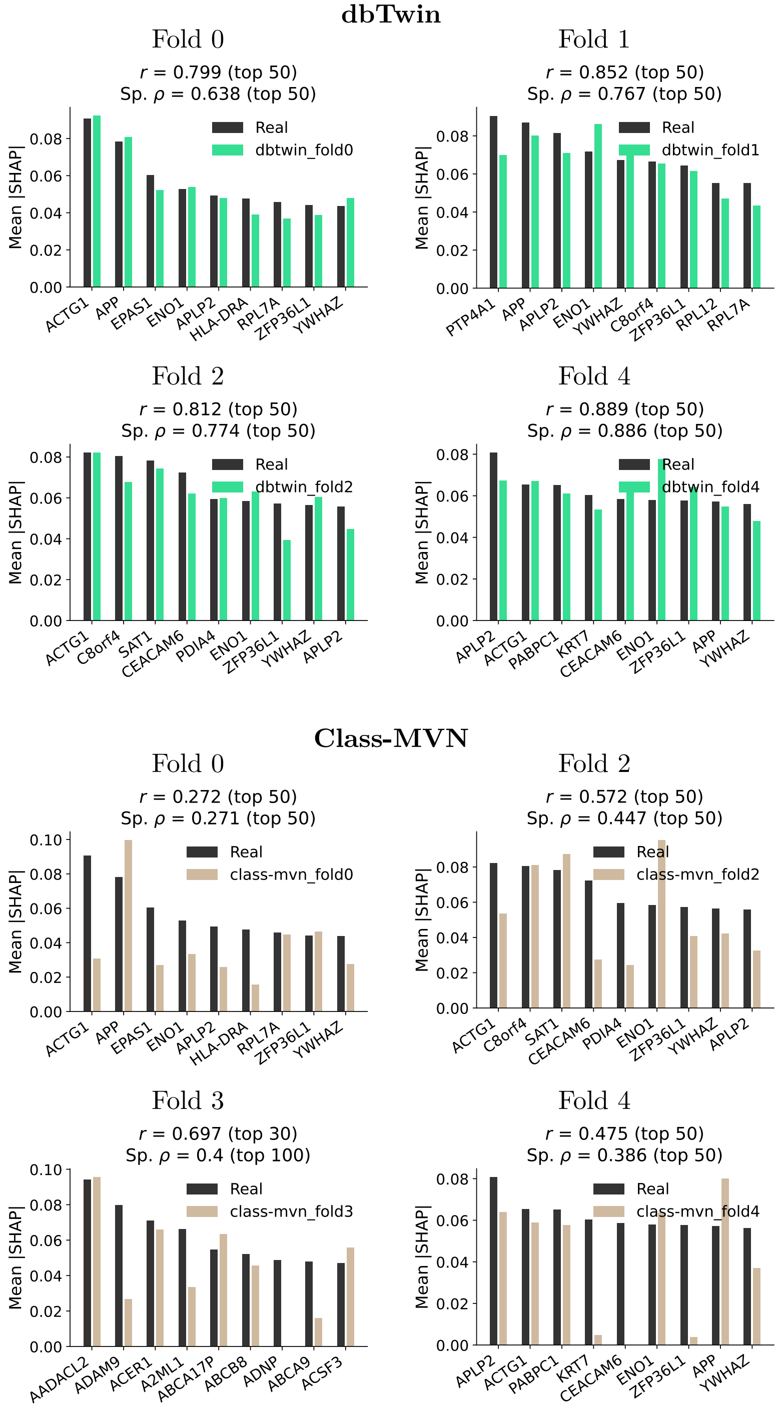


**Figure 5. SHAP gene-feature concordance for TCGA-LUAD EGFR mutation dataset in 4 remaining folds not shown in main text.** Top panel: **dbTwin**: SHAP plots for each fold. Bottom panel: **Class-MVN** SHAP plots for each fold.

### SHAP Analysis — RISK Pediatric Disease

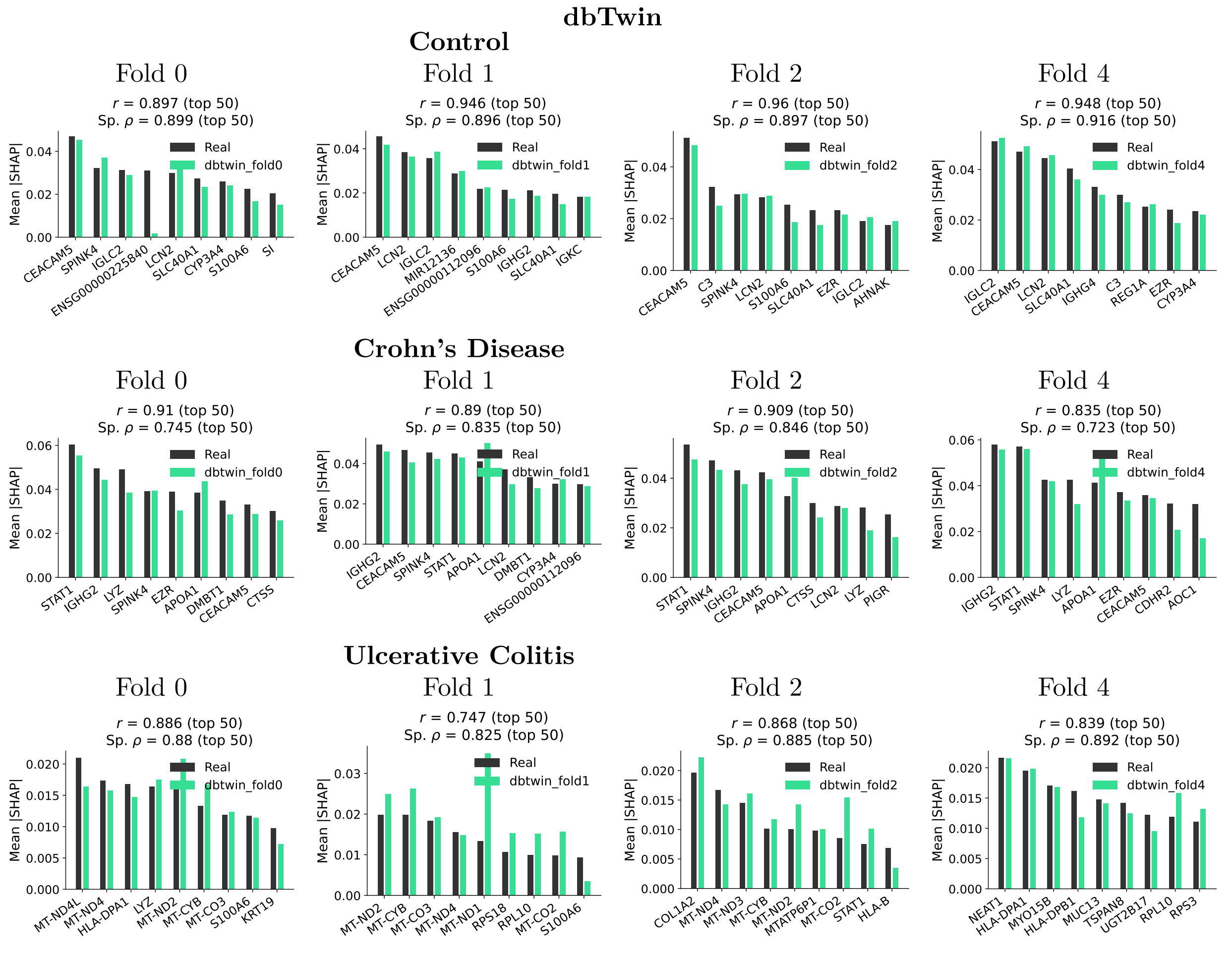


**Figure 6. dbTwin: SHAP gene-feature concordance for RISK pediatric disease dataset across 4-folds not shown in main text** Results organized by disease class (Control, Crohn's Disease, Ulcerative Colitis) with available folds.

### SHAP Analysis — RISK Pediatric Disease (Class-MVN)

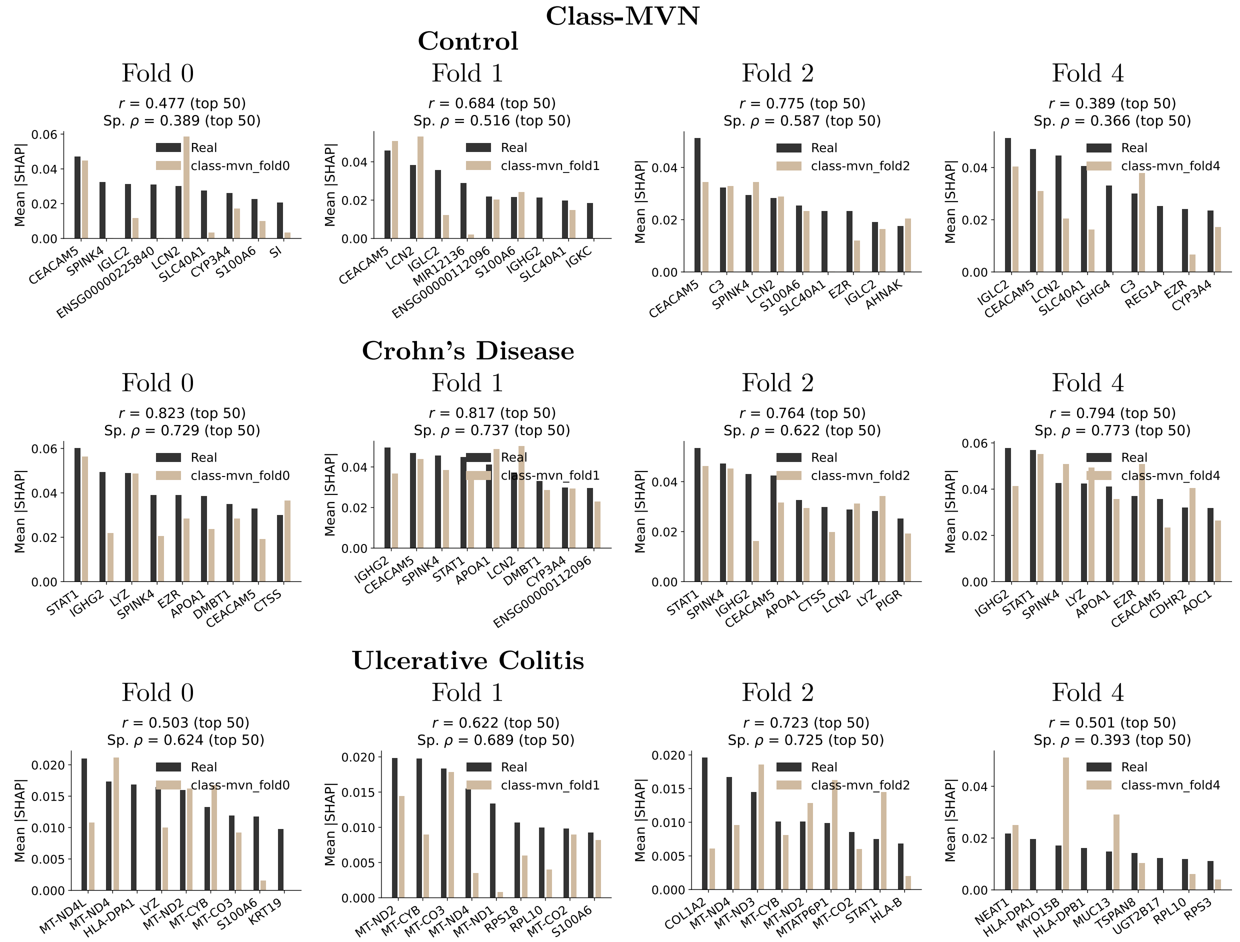


**Figure 7. class-MVN: SHAP gene-feature concordance for RISK pediatric disease dataset for four folds not shown in main text.** Results organized by disease class (Control, Crohn's Disease, Ulcerative Colitis) with available folds.

### SHAP Analysis — TCGA-LUAD Clinical Stage

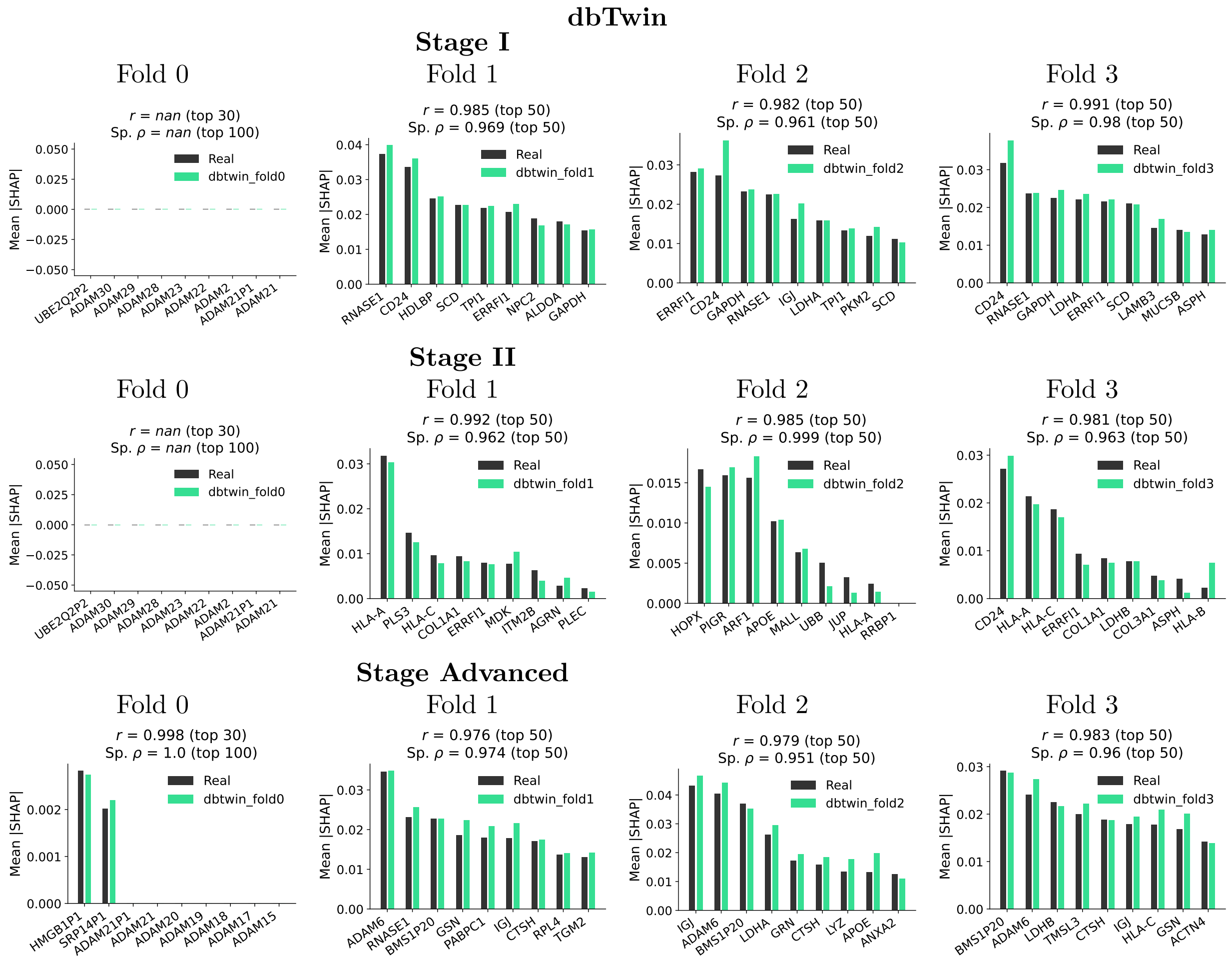


**Figure 8. dbTwin SHAP gene-feature concordance TCGA-LUAD clinical stage dataset for 4 folds not shown in main text.** Results organized by clinical stage (Stage I, Stage II, Stage Advanced).

### SHAP Analysis — TCGA-LUAD Clinical Stage (Class-MVN)

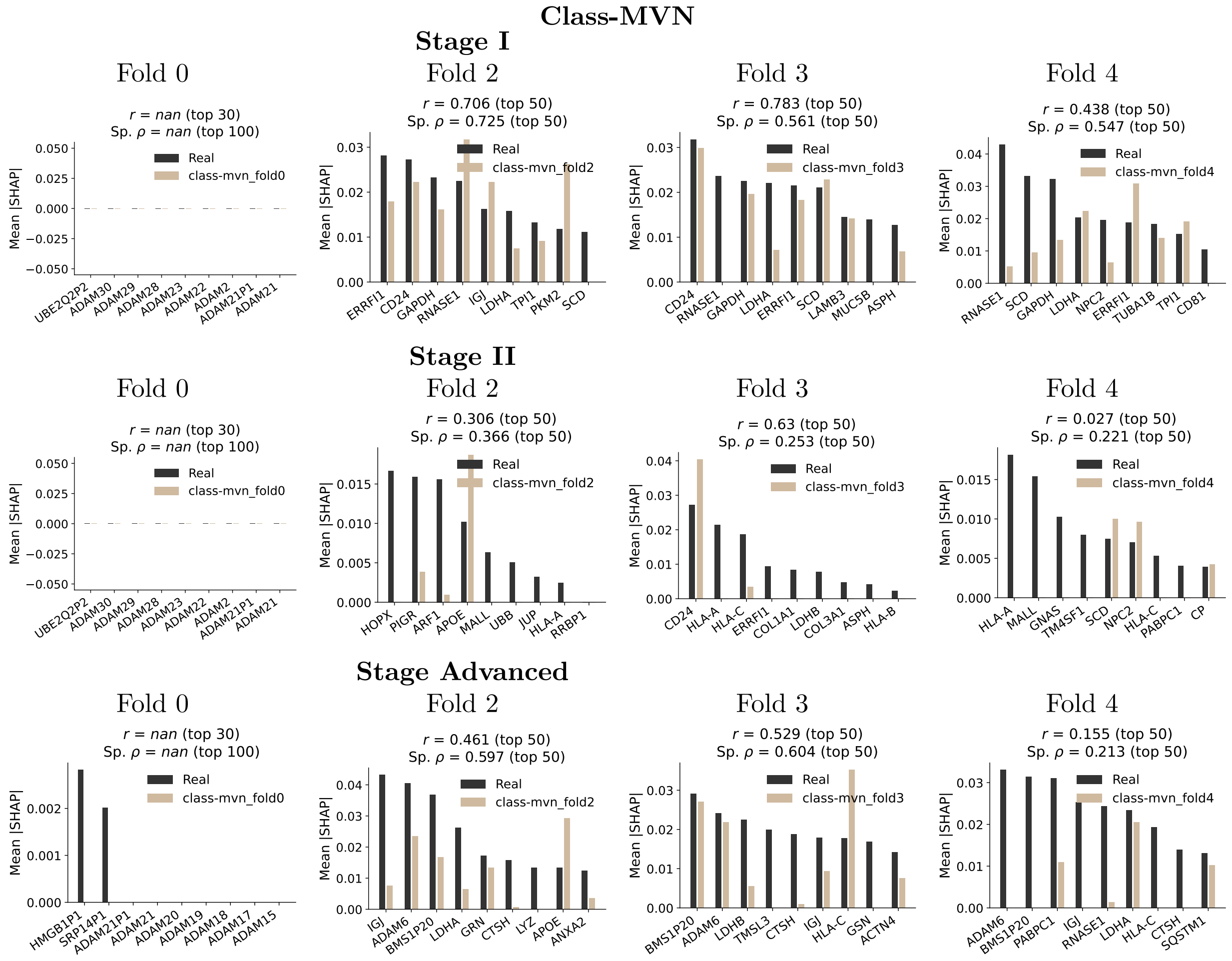


**Figure 9. class-MVN SHAP gene-feature concordance for TCGA-LUAD clinical stage dataset across 4 folds not in main text** Results organized by clinical stage (Stage I, Stage II, Stage Advanced) with available folds.

### Distance to Closest Record (DCR) — TCGA-LUAD EGFR

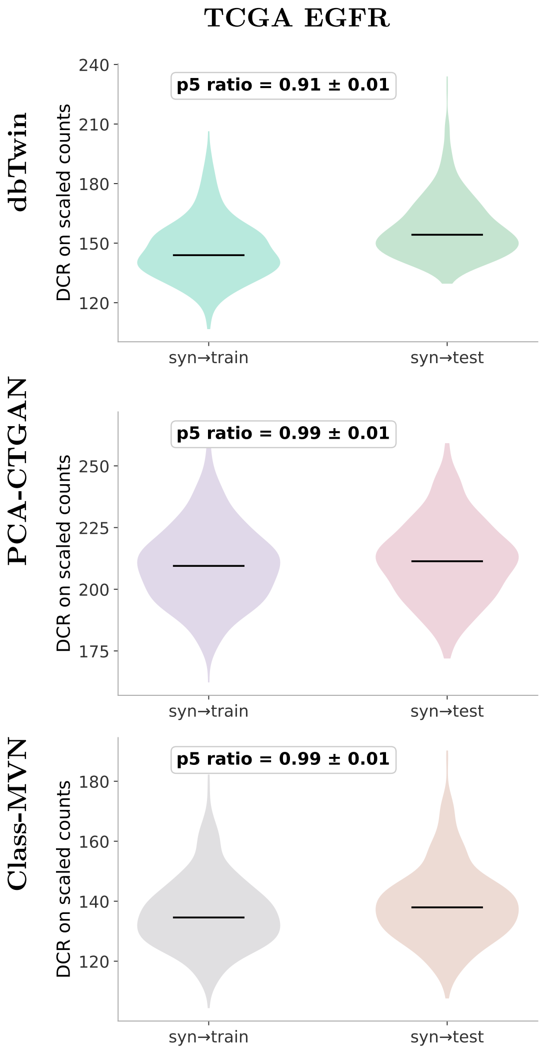


**Figure 10. Distance to closest record (DCR) privacy metric for LUAD-EGFR dataset.** Results shown for three generators (rows). DCR distributions compare the distance from each synthetic sample to its nearest neighbor in the real training set (blue) versus the held-out real test set (orange). Higher separation between distributions indicates lower re-identification risk.
